# *In silico*, biochemical, and *in vitro* analysis of silvestrol binding to the DEAD box RNA helicase eIF4A reveals broad anti-pathogen potential of rocaglates across the eukaryotic tree of life

**DOI:** 10.1101/2022.09.09.507249

**Authors:** Wiebke Obermann, Mohammad Farhan Darin Bin Azri, Leonie Konopka, Nina Schmidt, Francesca Magari, Julian Sherman, Liliana M. R. Silva, Carlos Hermosilla, Andreas H. Ludewig, Hicham Houhou, Simone Haeberlein, Mona Yiting Luo, Irina Häcker, Marc F. Schetelig, Christoph G. Grevelding, Frank C. Schroeder, Gilbert Lau, Anja Taubert, Ana Rodriguez, Andreas Heine, Tiong Chia Yeo, Arnold Grünweller, Gaspar Taroncher-Oldenburg

## Abstract

Selective inhibition of eukaryotic initiation factor 4A (eIF4A), an RNA helicase, has been proposed as a strategy to fight pathogens. Plant-derived rocaglates exhibit some of the highest specificities among eIF4A inhibitors. Sensitivity to rocaglates is determined by key amino acid (aa) residues mediating reversible clamping of the eIF4A:RNA complex. To date, no comprehensive assessment of eIF4A sensitivity to rocaglates across the eukaryotic tree of life has been performed to determine their anti-pathogenic potential.

We performed an *in silico* analysis of the substitution patterns of six aa residues in eIF4A1 critical to rocaglate binding (human positions 158, 159, 163, 192, 195, 199), uncovering 35 pattern variants among 365 eIF4As sequenced to date. *In silico* molecular docking analysis of the eIF4A:RNA:rocaglate complexes of the 35 variants, modeled in a human eIF4A environment, and *in vitro* thermal shift assays with recombinantly expressed human eIF4A mutants, representing select natural and artificial variants, revealed that sensitivity to a natural or one of two synthetic rocaglates—silvestrol, CR-1-31-B, or zotatifin—was associated with lower inferred binding energies and higher melting temperature shifts. Helicase activities were comparable across variants and independent of sensitivity to rocaglates.

*In vitro* testing with silvestrol validated predicted resistance based on position 163 substitutions in *Caenorhabditis elegans* and *Leishmania amazonensis* and predicted sensitivity in *Aedes sp.*, *Schistosoma mansoni*, *Trypanosoma brucei*, *Plasmodium falciparum*, and *Toxoplasma gondii*.

Our analysis shows resistance to rocaglates emerging in disparate eukaryotic clades pointing to resistance being a selective neutral trait except in rocaglate-producing *Aglaia* plants and their fungal parasite *Ophiocordyceps*. The analysis further revealed the possibility of targeting important insect, plant, animal, and human pathogens including *Galleria mellonella*, *Ustilago maydis*, *Babesia ovata*, and *Cryptosporidium sp.*, with rocaglates. Finally, combined docking and thermal shift analyses might help design novel synthetic rocaglate derivatives or alternative eIF4A inhibitors to fight pathogens.

**Author Summary:** In the ongoing search for novel ways to fight non-viral and non-bacterial pathogens, targeting translation—the universal process of protein synthesis—to inhibit growth and cell proliferation has emerged as an attractive strategy. Here, we focused on the potential of rocaglates, a group of plant-derived compounds, to inhibit an early step in translation mediated by a RNA helicase called eIF4A. We performed a comprehensive analysis of eIF4A sequence variants to determine their potential sensitivities to rocaglates, especially in pathogens of prokaryotic, fungal, or animal origin. We complemented this *in silico* analysis with enzyme-based *in vitro* and whole pathogen *in vivo* experiments to confirm the sensitivity or resistance to rocaglates of specific variants of eIF4A. Our analysis provides the first comprehensive picture of rocaglate sensitivity among pathogens and establishes targeting important insect, plant, animal, and human pathogens such as wax moth larvae, a major parasite of honey bees, corn smut, a widely distributed fungal disease, *Babesia*, a livestock parasite that causes anemia and babesiois, and *Cryptosporidium*, the causative organism of cryptosporidiosis in humans, with rocaglates as a viable anti-pathogen strategy.

## Introduction

Targeting eukaryotic translation has recently emerged as a potentially viable strategy to combat important pathogens such as the malaria-causing protozoan *Plasmodium falciparum*, the African trypanosomiasis–causing protozoan *Trypanosoma brucei*, the candidiasis-causing fungus *Candida auris*, or the lymphatic filariasis–causing nematode *Brugia malayi* [1-4]. Of the three steps that constitute translation—initiation, elongation and termination, initiation has garnered particular interest as a target for inhibiting translation in eukaryotic pathogens due to the diversity of factors involved in the process and its rate limiting effect on the overall translation process [5-7].

The eukaryotic translation initiation factor 4A (eIF4A), an ATP-dependent DEAD-box RNA helicase, is a highly conserved protein that plays an essential role in the initiation of translation and protein synthesis in eukaryotes [8]. Together with the cap-binding protein eIF4E and the scaffold protein eIF4G, eIF4A forms the eukaryotic translation initiation complex eIF4F. This complex is responsible for ‘activating’ and unwinding 5’-capped cellular mRNAs, making them accessible to the 40S ribosomal subunit and ultimately leading to the formation of the elongation-competent 80S complex and subsequent protein synthesis [9].

There are two isoforms of eIF4A in mammals that exhibit a sequence identity of 90- 95% and have equivalent biochemical functions: eIF4A1 and eIF4A2 [10,11]. Both isoforms differ significantly in expression levels *in vivo*, with eIF4A1 being present in almost all tissues during active cell growth and eIF4A2 mainly in organs with low proliferation rates [12]. A third ortholog, eIF4A3, shares only ∼60% sequence identity with eIF4A1 or eIF4A2 and is functionally distinct—eIF4A3 is involved in the assembly of the exon junction complex, which coordinates splicing of pre-mRNAs, but not in the formation of eIF4F [10].

Rocaglates, a class of plant-derived flavaglines containing a cyclopenta[*b*]benzofuran structure, are among the most potent and specific eIF4A inhibitors known to date [13, 14; **Figure S1**]. Over 200 natural and synthetic rocaglates have been described since rocaglamide A (RocA) was first isolated from Asian mahogany (*Aglaia sp*.) in 1982 [15-18]. Besides eIF4A, only two other molecular targets of rocaglates have been identified, albeit neither of them has been shown so far to be central to cell viability: the RNA helicase DEAD-box polypeptide 3 (DDX3), which facilitates translation of mRNAs with highly stable RNA secondary structures at the terminal part of the 5’UTR, and Prohibitins 1 and 2 (PHB1; PHB2), which as a complex contribute to the regulation of mitochondrial activity [19, 20].

Importantly, rocaglates preferentially clamp eIF4A:RNA complexes containing RNA strands with stable secondary structures such as stem-loops or G-quadruplexes and also polypurine stretches in the 5’UTR, all associated with subclasses of mRNAs such as proto-oncogenes or viral mRNAs [21-23]. mRNAs with complex secondary structures are often associated with proliferating cells, play a central role in translational regulation of many organisms, including pathogens, and are preferentially processed by eIF4A [24-26]. The preference of eIF4A to unwind mRNAs with stable secondary structures in their 5′UTRs, together with the specific avidity of rocaglates for clamping eIF4A:RNA complexes containing RNAs with stable secondary structures, makes this a potentially viable approach to fight pathogens because the resulting low toxicities for humans and animals—less is known about plants—could result in potentially favorable therapeutic windows [27].

The therapeutic potential of inhibiting eIF4A with rocaglates has already been shown *in vivo* in the context of cancer—the synthetic analog zotatifin is in a Phase 1/2 clinical study in patients with solid tumors [28]—and for host-targeted antiviral activity—zotatifin is in a dose escalating Phase 1 clinical study in patients with mild or moderate COVID-19 [29]. Indeed, multiple *in vitro* and *in vivo* studies have now shown the effectiveness of rocaglates in preventing the replication of RNA viruses including corona-, picorna-, flavi-, filo-, hepe-, toga-, arena-, nairo, and bunya viruses, supporting the potential of rocaglates as pan-antivirals for minimizing the impact of future RNA virus pandemics [30-35].

Several reports have further shown the eIF4A-dependent anti-pathogenic potential of rocaglates against *Plasmodium falciparum* and *P. berghei*, *Candida auris*, and a number of other eukaryotic pathogens [36, 37; see **Table S1**]. Other pathogens have also been shown to be resistant to rocaglates, including *Entamoeba histolytica* and *Leishmania donovani* [38, 39; see **Table S1**]. To date, however, no comprehensive analysis has been performed to assess the true anti-pathogenic potential of rocaglates.

Rocaglates reversibly bind to eIF4A:RNA complexes, leading to a stable ternary complex that prevents enzymatic unwinding of the secondary structure of the mRNA [40, 41]. Select aa within the rocaglate/RNA-binding pocket—human eIF4A1 position 158, 159, 163, 192, 195, 199—have been shown to be critical to the rocaglate clamping mechanism and to determine resistance versus sensitivity to rocaglates [41-43]. Initial studies in yeast have shown how single substitutions in all six aa could cause resistance to silvestrol and the synthetic rocaglamide ROC-N without affecting the helicase activity of eIF4A [42]. *In vitro* and *in vivo* mouse studies confirmed the critical roles of substitutions at positions 163 and 199 (for the remainder of this article we will be referring to the human position numbering) [43]. Subsequently, analysis of the crystal structure of human eIF4A1 in complex with RocA, polypurine RNA, and an non-hydrolyzable ATP analogue, has revealed that rocaglates reversibly clamp mRNAs to eIF4A through π-π-stacking interactions between the rocaglates’ A and B phenyl rings and two consecutive purines in the bound RNA substrate, and between the rocaglates’ C phenyl ring and the amino acid residue at position 163, confirming the previous *in vitro* and *in vivo* studies [41]. Incidentally, the eIF4A sequences of rocaglate- producing *Aglaia* spp. exhibit resistant substitutions at both aa 163 (Phe to Leu) and aa 199 (Ile to Met), and a recent report has revealed the presence of another resistance conferring substitution at aa 163 (Phe to Gly) in *Ophiocordyceps* sp. BRM1, a fungal parasite of *Aglaia* [41, 44].

We rationalized that a comprehensive *in silico* analysis of published eIF4A sequences, together with a biochemical analysis of representative sequences and *in vitro* studies with eukaryotic microorganisms containing previously untested variants of eIF4A, would provide a much more complete picture of the anti-pathogenic potential of rocaglates, shed light on the potential evolutionary significance of rocaglate resistance, and further our insights into structure-activity relationships that could inform the design of next-generation rocaglate derivatives or other eIF4A inhibitors for anti-pathogen use.

## Results

### Global eIF4A sequence analysis reveals limited diversity of rocaglate- interacting aa patterns in the RNA-binding pocket

We performed a global GenBank (https://www.ncbi.nlm.nih.gov/genbank/) search for published eIF4A sequences that resulted in the identification of 365 unique eIF4A1 and eIF4A2 protein sequences—78 protist, 80 fungal, 49 plant, and 158 animal (**Table S2**). No eIF4A3 were included in our analysis. Of the 365 sequences, 162 corresponded to pathogens of protist (55), fungal (53), or animal (54) origin. To determine the potential interactions of the different eIF4A proteins with rocaglates, we analyzed the substitution patterns of the six aa residues located between motifs Ib and III of eIF4A that have been previously identified as being critical to rocaglate binding (human positions 158, 159, 163, 192, 195, 199) [41, 42] (**Figure S2**). These aa residues are located in and around the eIF4A RNA-binding pocket and include three residues directly involved in RNA binding (159, 192, and 195) [45]. This region of eIF4A also includes additional RNA-binding aa residues (160, 161) and two residues involved in interactions with ATP (182, 183) [46, 47]. The aa patterns known to date to be associated with sensitivity or resistance to rocaglates are listed in **Table S1**.

Our analysis uncovered 35 aa patterns, with just four of them—T158, P159, Y163, F192, Q195, V199; TPFFQI; TPFFQV; TPYFQI—accounting for 63% of all known eIF4A sequences and represented in all four major groups of eukaryotes—protists, fungi, plants, and animals (**Figure S3**). On the other end of the spectrum, over half of the aa patterns (24/35) were seen in only one of the groups of eukaryotes. Of note are the patterns TPLFQM, exclusively present in all rocaglate-producing *Aglaia* spp., including the two species reported here, *Aglaia stellatopilosa* and *Aglaia glabriflora* (**Figure S2**; Accession numbers **XXX** and **XXX**, respectively), and TPGFQI, reported in only one species of the fungal genus *Ophiocordyceps* that was identified as a parasite of *Aglaia* spp..

Among the 162 pathogens analyzed, we identified 24 different aa patterns with the protists showing the highest diversity (14) followed by fungi (13) and animals (9) (**Table S2**). Five of the patterns—TPYFQV, TPFFQI, TPLFQI, TPHFQV, TPFFQV—accounted for 53% of all pathogen eIF4A sequences and were represented in protists, fungi, and animals (**Figure 1**). Conversely, two thirds of the aa patterns (16/24) were seen in only one of the groups of eukaryotes. All four aa patterns known to confer sensitivity to rocaglates—TPYFQV, TPFFQI, TPYFQI, TPHFQI—could be detected among the pathogens analyzed, representing 50% of the sequences (**Figure 1**). Another 14% of the sequences (22/162) exhibited aa patterns—TPLFQI, TPSFQI, TPGFQI—previously associated with resistance to rocaglates, suggesting that resistance among pathogens is low and mostly present among protists (**Table S1**).

**Figure 1:**
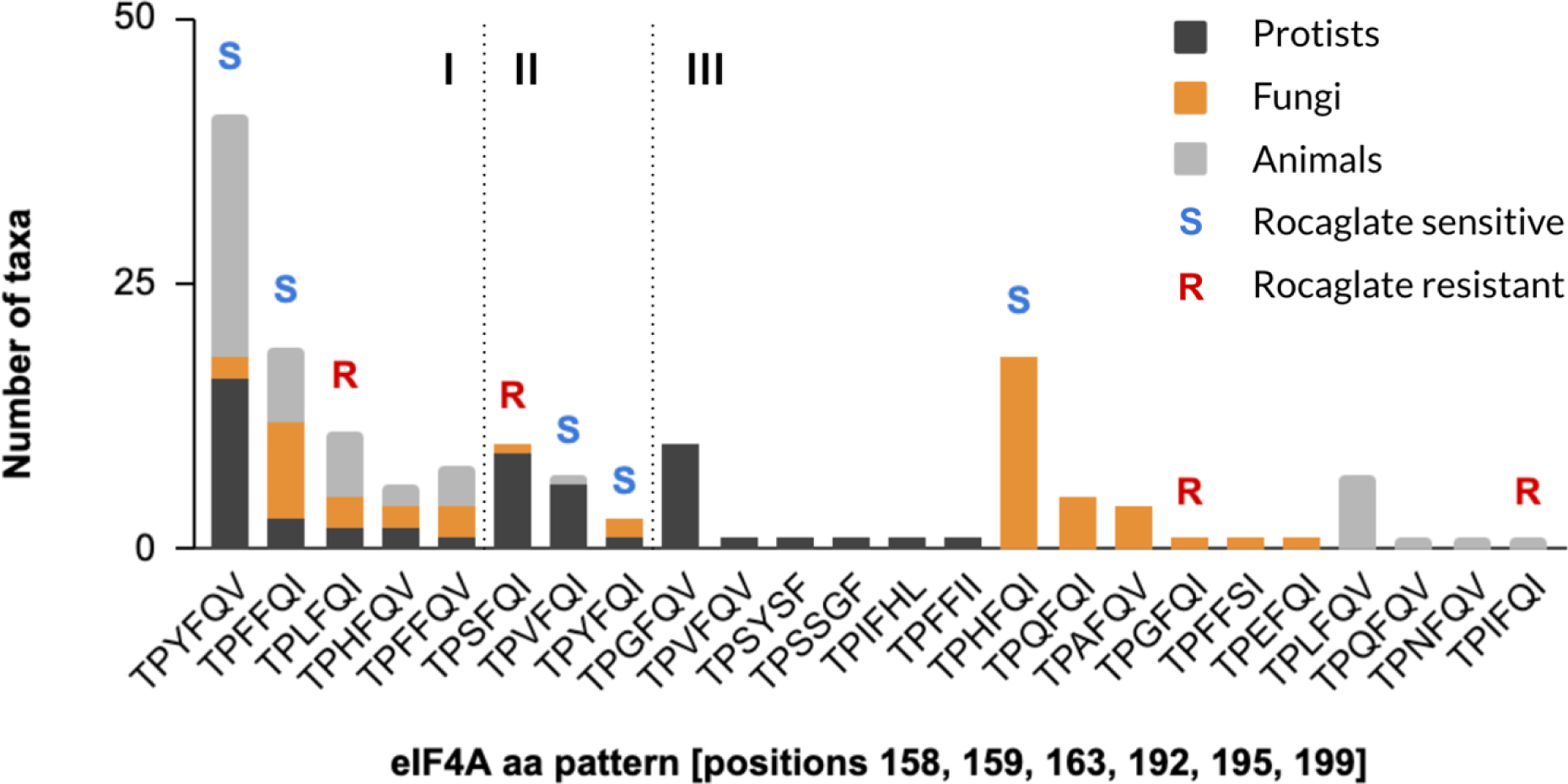
Representation of patterns of aa critical for rocaglate-binding in known eIF4A proteins across the three main groups of eukaryotic pathogens. A comprehensive analysis of known eIF4A proteins encoded by pathogens revealed 24 patterns of aa at positions 158, 159, 163, 192, 195, and 199 (human eIF4A1 numbering). Five aa patterns were present in all three groups of eukaryotes (I), three patterns were present in two groups (II) and sixteen patterns were present in only one group (III). Patterns known to provide natural sensitivity or resistance to rocaglates could be found in all three groups.

Pathogens we identified as potentially sensitive to rocaglates include the parasitic roundworm *Trichuris trichiura*, which causes helminthiasis in humans, the tsetse fly *Glossina morsitans*, a major vector of the African trypanosomiasis–causing parasite *Trypanosoma brucei*, the fungus *Aspergillus niger*, which causes black mold in fruits and vegetables, and the parasite *Cryptosporidium* sp., the causative organism of cryptosporidiosis (**Table S2**). Among the pathogens potentially resistant to rocaglates we identified the gastrointestinal and lung parasite *Paragonimus westermani*, the causative agent of lung fluke disease, the plant pathogen *Blumeria graminis*, which causes powdery mildew on cereals, and *Leishmania* sp., a genus of protozoan parasites that causes leishmaniasis.

### Substitution tolerance analysis of rocaglate-interacting aa patterns reveals stark dichotomy between highly conserved and highly variable residues

To ascertain the natural tolerance for substitutions at the six rocaglate-interacting positions of eIF4A and their potential evolutionary implications, we quantified the corresponding aa distributions across all 365 sequences evaluated in this study. Substitution tolerance at the six rocaglate-binding positions were markedly different, with four positions—158, 159, 192, 195— being highly conserved and the two other positions—163, 199—showing different degrees of variation (**Figure 2**). Amino acid positions 158, 192, and 195 are involved in RNA binding, and position 159, while itself not directly involved in RNA binding, is flanked by three RNA-binding residues—158, 160, and 161 (**Figure S2**) [45]. By contrast, aa position 163, which together with positions 158 and 159 is part of the highly conserved motif Ib of DEAD-box RNA helicases and is directly adjacent to another RNA-interacting residue at position 164, showed a high level of tolerance for substitutions, with 14 different aa filling the position across all eIF4As we surveyed. The six aa not detected in position 163 fall within different biochemical categories—positively-charged basic [Arg, Lys], nonpolar aliphatic [Met, Pro], nonpolar aromatic [Trp], and polar [Thr]—and have been associated with destabilization of α-helices in proteins [Pro] or modulation of protein interactions with nucleic acids [Arg] among others [48- 50]. Finally, position 199 exhibited a quasi bimodal distribution with an almost equal number of sequences encoding an Ile or a Val. Position 199 is part of an α-helix situated between motifs II and III of eIF4A, which might explain the tolerance for Ile or Val, two structurally similar aliphatic aa of equivalent hydrophobicity. Interestingly, at the codon level, the switch from Ile to Val only requires the change of the first nucleotide from adenine to guanine (AUA, AUU, or AUC to GUA, GUU, or GUC), and the switch from Ile to Met, only observed in *Aglaia*, requires a change of the third nucleotide, the ‘wobble position’, in any of the Ile codons to guanine (AUA, AUU, or AUC to AUG) [51]. The same applies to the switch from Phe (UUC or UUU) to Leu (UUG or UUA).

**Figure 2:**
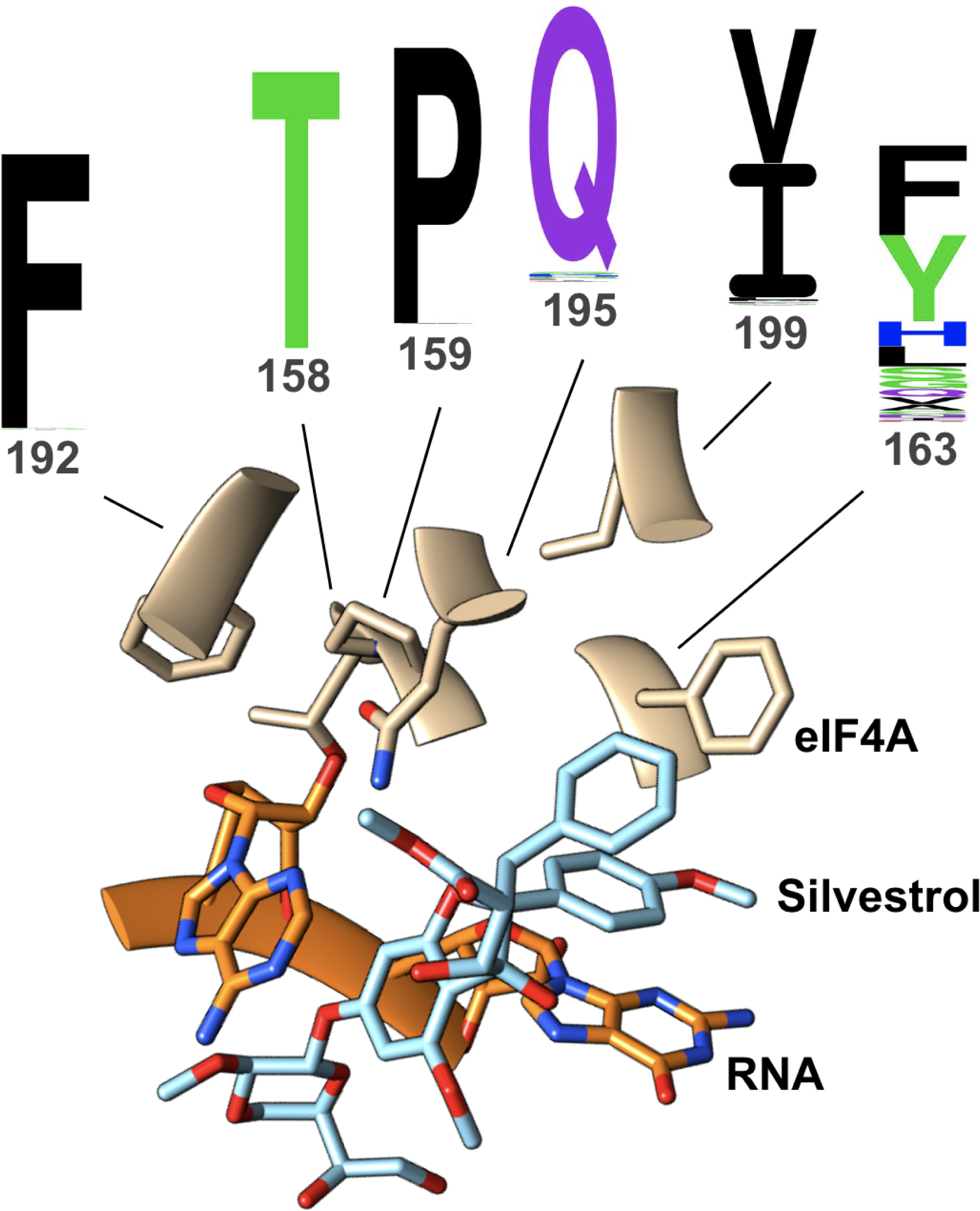
Amino acid substitution profiles at six positions known to be critical for rocaglate binding. Six aa residues—158, 159, 163, 192, 195, 199—in the eIF4A RNA-binding pocket have been shown to be critical for rocaglate binding. Natural substitution tolerances at each of the six residues across the 365 eIF4A sequences analyzed here revealed four highly conserved positions—158, 159, 192, 195—and two variable positions—163 and 199. Of these, the latter shows a quasi bimodal frequency distribution (I or V), while the former shows a more promiscuous frequency distribution, with 14 different aa being tolerated in that position. The structure depicted above corresponds to the aa pattern TPFFQI; the rocaglate shown interacting with the eIF4A:RNA complex is silvestrol. The sequence logos were generated with Seq2Logo - 2.0 [53].

The differential natural substitution tolerance profiles we determined confirmed previous reports that had suggested position 163 and, to a lesser extent, position 199 of the eIF4A protein were the two key residues to evaluate when determining potential resistance to rocaglates [41, 42]. The strict conservation of residues T158, P159, F192, and Q195, together with the seemingly neutral flip between Val and Ile at position 199, point to the residue at position 163 as the key determinant of sensitivity to rocaglates in eIF4A. Incidentally, several of the aa substitutions we observed at position 163—Glu, Asp, His, Phe, Tyr—have been reported to have the capacity to establish either π-π-stacking or hydrophobic interactions and thus render the variants sensitive to rocaglates [52].

Finally, the substitution pattern at position 163 also seemed to indicate ‘evolutionary preferences’ across the four groups of eukaryotes. In protists, 29% of the sequences had Tyr in position 163, followed by Ser (15%), Phe (14%), Gly (14%), and Val (12%). In fungi the distribution was His (36%), Phe (24%), Gln (12%), and Ala (10%). In plants, 73% of the eIF4A sequences had Phe in position 163 and 10% Tyr, while in animals the proportion was switched to 51% Tyr and 34% Phe (**Table S2**; **Figure S3**).

### Evolutionary analysis of eIF4A aa patterns associated with rocaglate sensitivity suggests emergence of resistance is serendipitous

To gain a better understanding of the evolutionary context for the emergence of resistance to rocaglates and its potential implications for the use of rocaglates as anti-pathogens, we projected the 365 sequences we analyzed onto the latest version of the eukaryotic tree of life (eToL) (**Figure 3**) [54]. Out of the eleven clades represented in the eIF4A sequence analysis, six, including deep-rooted branches such as the discoba and metamonada, contained organisms resistant to rocaglates—*Leishmania* sp. among discoba and *Giardia* sp. among metamonada.

**Figure 3:**
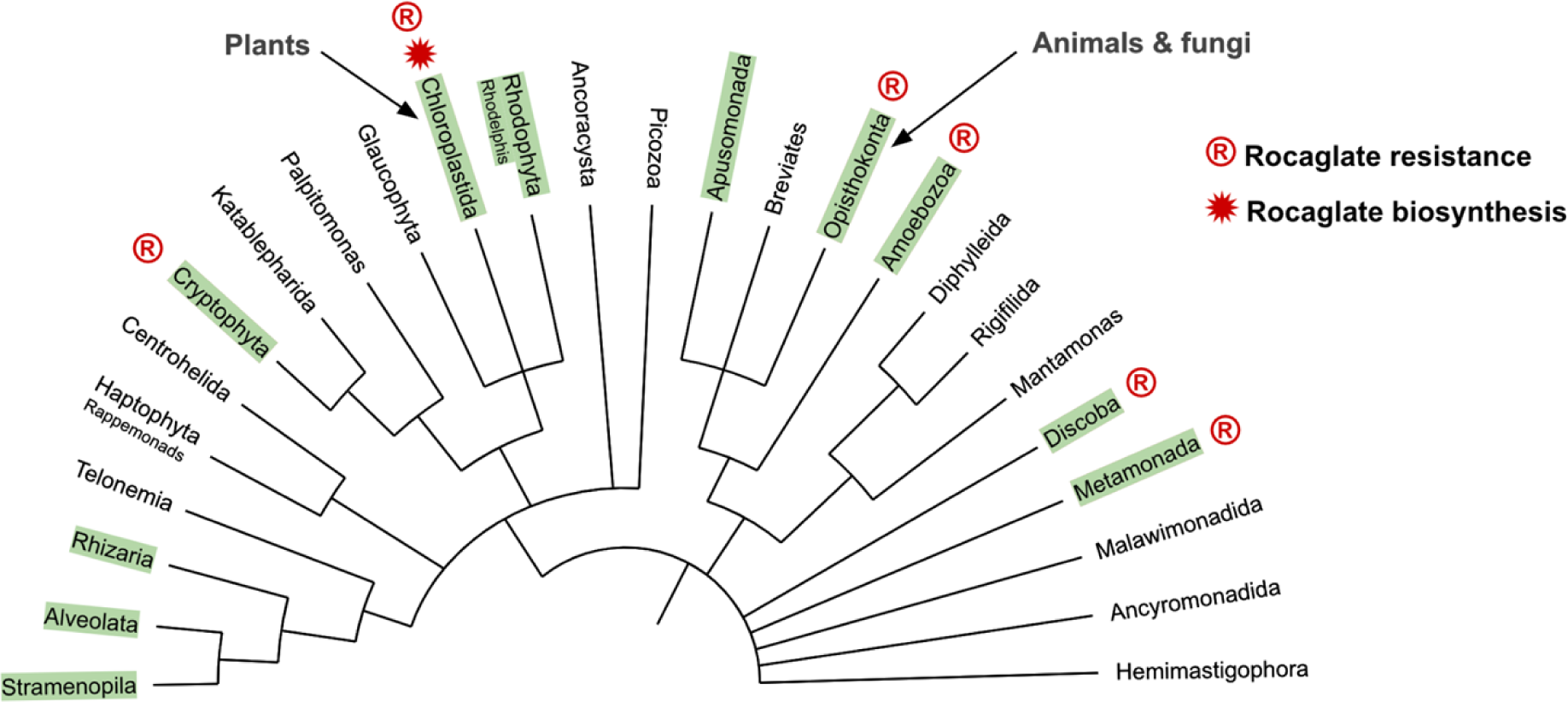
Distribution of eIF4A rocaglate resistance on the eToL. Green boxes denote clades represented in our eIF4A analysis. Rocaglate resistant variants can be found in deep rooted as well as more recently evolved clades. Several variants are exclusive to one clade while others are distributed more generally.

Intriguingly, the only organisms known to produce rocaglates are plants within the genus *Aglaia*, which like all higher plants emerged much later in evolution. This asynchrony in the emergence of rocaglate synthesis and eIF4A resistance to it indicates that the emergence of ‘rocaglate resistance’ would not have been driven by the evolutionary pressure of exposure to rocaglates but rather occurred as a serendipitous outcome of natural diversification of the eIF4A sequence wholly unrelated to rocaglates. The single exception to this rule would be *Ophiocordyceps* sp. BRM1, the *Aglaia* sp. parasitic fungus, which might have developed resistance to rocaglates through direct exposure [44].

The recent determination of the crystal structure of the eIF4A:RNA complex in association with Pateamine A (PatA), a macrolide isolated from the marine sponge *Mycale hentscheli*, provides incidental support for this evolutionary conjecture [55, 56]. Despite having a completely unrelated structure, PatA interacts with the eIF4A:RNA complex in an analogous way to rocaglates, and this functional mimicry extends to the inability of PatA to stabilize the eIF4A:RNA complex in the presence of an F163L substitution [55, 57]. Similarly to rocaglates, PatA is biosynthesized by an organism, *Mycale hentscheli*, that is deeply rooted in the animal tree, one of the most recent branches of the Opisthokonta. The protein sequence of the *Mycale hentscheli* eIF4A has not been determined, and no comprehensive map of eIF4A resistance/sensitivity to PatA exists, but the fact that resistance to this natural product is present in early, non-opisthokonta lineages that were presumably never exposed to PatA over evolutionary timescales, is intriguingly similar to the scenario depicted above for rocaglates.

In *Aglaia* itself, by contrast, the emergence of resistance would have been a necessary step happening in parallel or already in place as the ability to synthesize rocaglates evolved. The rocaglate-resistant aa pattern of the *Aglaia* eIF4A, TPLFQM, contains the most common ‘resistant’ substitution at position 163, Leu, and a Met substitution at position 199. It would be interesting to next sequence eIF4A in *Mycale hentscheli* to determine whether it also contains resistant substitutions and which.

### Subgroup of organisms contains eIF4A isoforms with divergent rocaglate- associated aa patterns

There are two isoforms of eIF4A in mammals that exhibit a sequence identity of 90-95% and have equivalent biochemical functions: eIF4A1 and eIF4A2 [10, 11]. Both isoforms differ significantly in expression levels *in vivo*, with eIF4A1 being present in almost all tissues during active cell growth and eIF4A2 mainly in organs with low proliferation rates [12]. Given the high sequence identity between both isoforms and the location of the aa residues involved in rocaglate interactions in the highly conserved RNA-binding pocket, no differences in the rocaglate-associated aa pattern have been reported.

Our global eIF4A sequence survey has now uncovered a number of organisms whose eIF4A isoforms contain divergent rocaglate-associated aa patterns (**Table S3**). Across the four groups of eukaryotes we analyzed, we uncovered several variations that fell into distinct categories. Among protists there were no divergent patterns except between the two eIF4A isoforms of the apusamonad *Thecamonas trahens*, which had the archetypal rocaglate-sensitive TPFFQI pattern on one isoform and a pattern containing three substitutions, TPLFAV, on the other. The Leu at position 163 points toward potential resistance to rocaglates, but no *in vitro* or *in vivo* confirmation exists. The fungal group contained five species exhibiting divergent isoforms. In all instances, the isoforms switched between a His and an Ala at position 163, accompanied in four cases with a concurrent switch from an Ile to Val in position 199; the fifth pair had a Val at this position in both isoforms. The five species belonged to five different genera, and three of them—*Neurospora crassa*, *Purpureocillium lilacinum*, and *Diplocarpon rosae*—are known pathogens. The effect of the H163A substitution on resistance to rocaglates has not been determined. Among plants, the six genera with divergent aa patterns exhibited a more diverse range of substitutions. Three of the isoform pairs switched between a Cys and a Phe at position 163—requiring just a point mutation of the second nucleotide of the corresponding codon from guanine to uracil (UGC or UGU to UUC or UUU)—accompanied by a concurrent switch from a Val to Ile in position 199. Another isoform pair switched between Ile and Val at position 199 but retained the Phe at position 163 in both isoforms, and the microalga *Raphidocelis subcapitata* had a pair of isoforms with a unique substitution pattern: Y163N and L199C. Finally, all species of rocaglate-producing *Aglaia* exhibited a unique combination of resistant isoforms, TPLFQM and TPLFQI.

We found the largest number of eIF4A isoforms containing divergent rocaglate- associated aa patterns among animal species. Of the 14 divergent isoform pairs we recorded, 11 exhibited likely neutral substitutions—F163Y and I199V—and five had an F163L substitution that would render one of the isoforms resistant and one sensitive to rocaglates. Incidentally, all diverging aa patterns with a resistant/sensitive dichotomy were found in species considered pathogenic—*Trichuris trichiura*, *Schistocephalus solidus*, *Hymenolepis microstoma*, and *Schistosoma mansoni*.

### Comparative analysis of RNA helicase activities of representative eIF4A variants suggests evolutionary convergence toward optimal enzyme performance independent of rocaglate resistance

The overlap between residues critical to the interaction of eIF4A with rocaglates and with RNA, raises the question of how particular substitutions at rocaglate-associated positions could affect the protein’s RNA helicase activity. A previous report indicates single mutations in positions 159, 163, and 195 have no or only little effect on RNA helicase activity [42]. To systematically determine the effect of different aa substitutions on the eIF4A RNA helicase activity, we generated 17 mutant eIF4A proteins with single and double substitutions at positions 163 and 199 in a human eIF4A1 background (**Table S4**). In addition to mutants recapitulating natural aa substitution patterns, we generated several variants containing non-natural aa changes to test potential evolutionary constraints on the nature of aa substitutions tolerated within the RNA-binding pocket.

Side-by-side evaluation of the RNA helicase activity profiles of the 17 eIF4A variants and the wild type human protein containing a TPFFQI aa pattern revealed a tight distribution of *V_max_* across variants recapitulating natural aa patterns (**Figure 4A**). The group of six non-natural substitution patterns showed a similar range of *V_max_* values to those measured for the natural patterns. Interestingly, three of the natural substitution patterns—two of them among the most highly represented in our sequence survey (TPFFQV; TPYFQI) and the third one being uniquely present in rocaglate-producing *Aglaia* sp. (TPLFQM)—exhibited the highest helicase activity. Among the non-natural aa patterns, a mutant derived from the *Aglaia* sp. pattern and containing an F163H and an I199M substitution also stood out in the non-natural variant group for exhibiting a *V_max_* more similar to that of these three high *V_max_* natural aa patterns.

**Figure 4:**
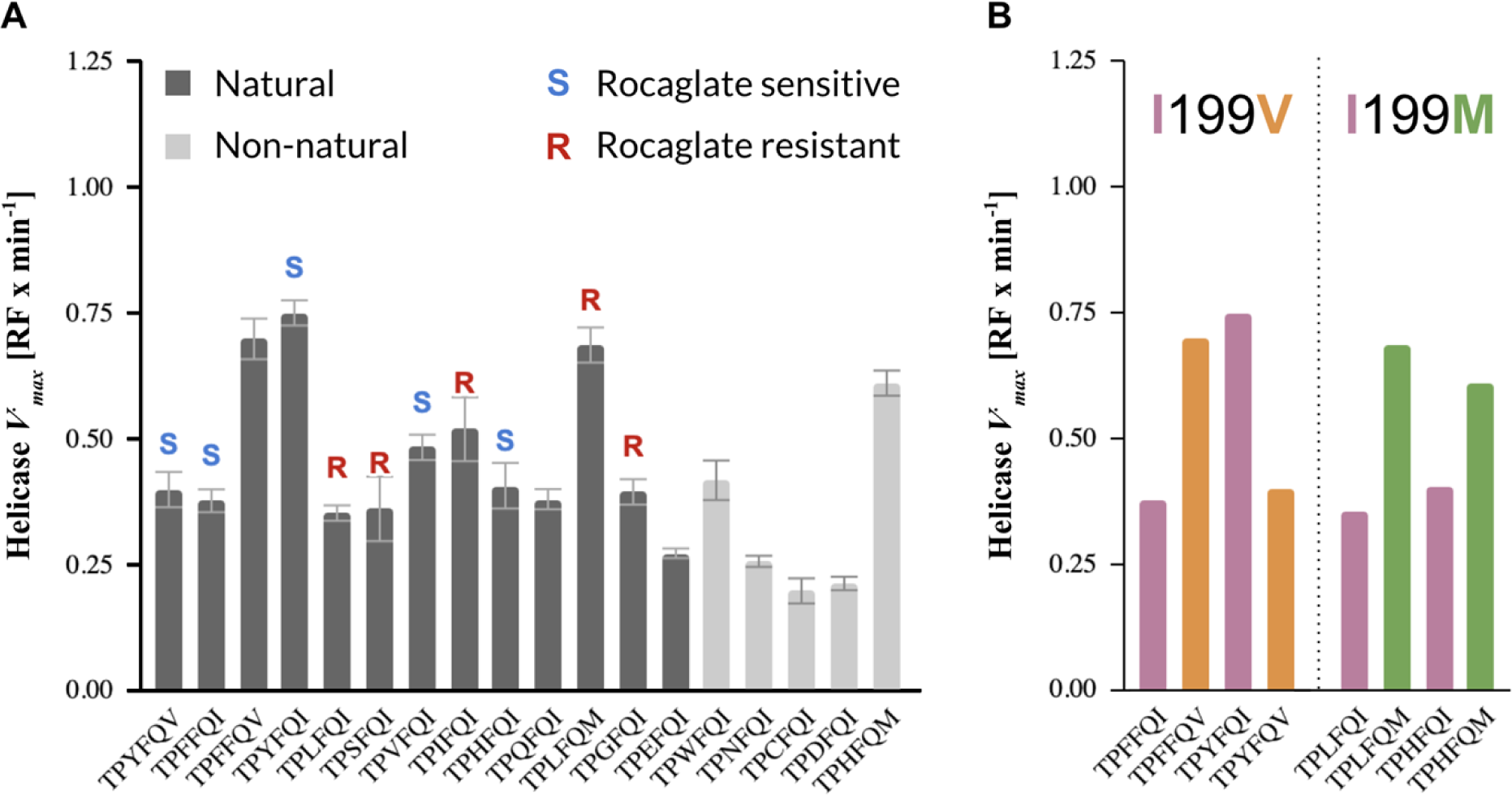
RNA helicase activities of eIF4A mutant proteins containing select natural and non-natural rocaglate-associated aa patterns in a human whole protein background. (**A**) Helicase *V_max_* of the wild-type human eIF4A1 containing the rocaglate-binding pattern TPFFQI and 17 variants of the human eIF4A1 protein (**Table S4**), including one expressing the *Aglaia* sp. aa pattern TPLFQM. All natural and non-natural aa patterns analyzed exhibited *V_max_* within a narrow range, indicating that the helicase activity is maintained. (**B**) Side-by-side *V_max_*comparison of mutant pairs differing only in the aa residue at position 199 of the eIF4A protein. The two substitutions analyzed, I199V and I199M, are the only natural substitutions at this position revealed by our global eIF4A sequence survey. (RF: relative fluorescence; error bars indicate mean standard error from three technical replicates; error bars removed in (**B**) for clarity)

A detailed look at two naturally occurring substitutions at position 199 of eIF4A— I199V and I199M—showed a modulating effect of these changes on the helicase activity (**Figure 4B**). The switch from Ile to Val enhanced or reduced *V_max_* depending on the aa at position 163. By contrast, in the two I199M mutants we analyzed, the switch from Ile to Met resulted in an equivalent enhancement of the corresponding *V_max_*, regardless of the aa at position 163. The similarity in effect size, directionality, or both, of these select single point mutations at position 199 on the *V_max_*, and the fact that the effect size lies squarely within the range of natural *V_max_* determined for natural eIF4A variants, indicate that the substitution tolerance at this position is determined by the efficiency of the helicase. With the exception of the *Aglaia sp.* eIF4A sequences, which contain a Met at position 199, the only two other residues tolerated at this position according to our survey are Ile and Val.

Similarly, the tolerance for substitutions at position 163 seemed to reflect a structural flexibility to accommodate a variety of aa residues of diverse characteristics without substantially affecting the helicase activity, and regardless of sensitivity to rocaglates. Importantly, all variants analyzed here were evaluated within a human whole protein scaffold. Even though eIF4A is a highly conserved protein, subtle differences in aa sequence throughout the length of the protein could affect folding and hence the ultimate structure of the RNA-binding pocket. The value of our analysis resides in the fact that all the substitutions were analyzed precisely within the same protein framework, allowing for the removal of other confounders. Ultimately, each of the patterns would have to be evaluated within its natural scaffold to determine the real *V_max_* of each eIF4A variant.

### Comparative thermal shift analysis of eIF4A:RNA:rocaglate complexes reveals direct correlation between rocaglate sensitivity and complex stability

The stability of the eIF4A:RNA:rocaglate complex is determined by the interactions between its three components. In particular, the clamping of the mRNA strand to eIF4A requires the establishment of π-π-stacking interactions, a process that is driven by the steric constraints of the mRNA-binding pocket. Natural eIF4A variants containing a Phe, Tyr, or His residue in position 163 have been reported to be sensitive to rocaglates, while natural variants with Leu or Ser substitutions have been shown to be resistant (**Table S2**). Based on the published structure of the eIF4A:RNA:RocA complex, Phe, Tyr, and His facilitate the establishment of π-π-stacking interactions while Leu and Ser do not provide a favorable molecular context for such interactions [41]. We set out to determine how other aa substitutions at position 163 would affect the stability of the π-π-stacking interactions by measuring the shifts in thermal denaturation temperature of different eIF4A:RNA:rocaglate complexes.

We analyzed 18 variants, including natural and non-natural variants, known to be resistant or sensitive to rocaglates, or for which no data on resistance were available (**Table S4**). The variants were chosen based on the natural prevalence of particular aa substitutions, e.g., Phe, Tyr, Leu, or His, and/or their potential for establishing π-π-stacking interactions, e.g., Trp. To evaluate the stabilizing effect of natural mutations in position 199, we designed the variants to alternate between Ile and Val as well as with Met, the characteristic aa substitution at this residue found in all *Aglaia sp.* eIF4As.

Thermal denaturation shifts, ***Δ*** melting temperature, were determined by comparing the melting temperatures of the eIF4A:RNA complex in the presence or absence of three different rocaglates: silvestrol, zotatifin, and CR-1-31-B. The ***Δ*** melting temperature measured for silvestrol showed a clear correlation between sensitivity to rocaglates and a larger shift in the thermal denaturation temperature (**Figure 5**). The larger shifts observed with the sensitive patterns point to the establishment of more stable eIF4A:RNA:rocaglate complexes that require more energy to dissociate due to the presence of stable π-π-stacking interactions. The amino acid patterns exhibiting the largest temperature shifts also happen to be the most well represented in nature within and across clades (**Figure 5**). We observed equivalent thermal denaturation shift patterns for zotatifin and CR-1-31-B (**Figure S4**).

**Figure 5:**
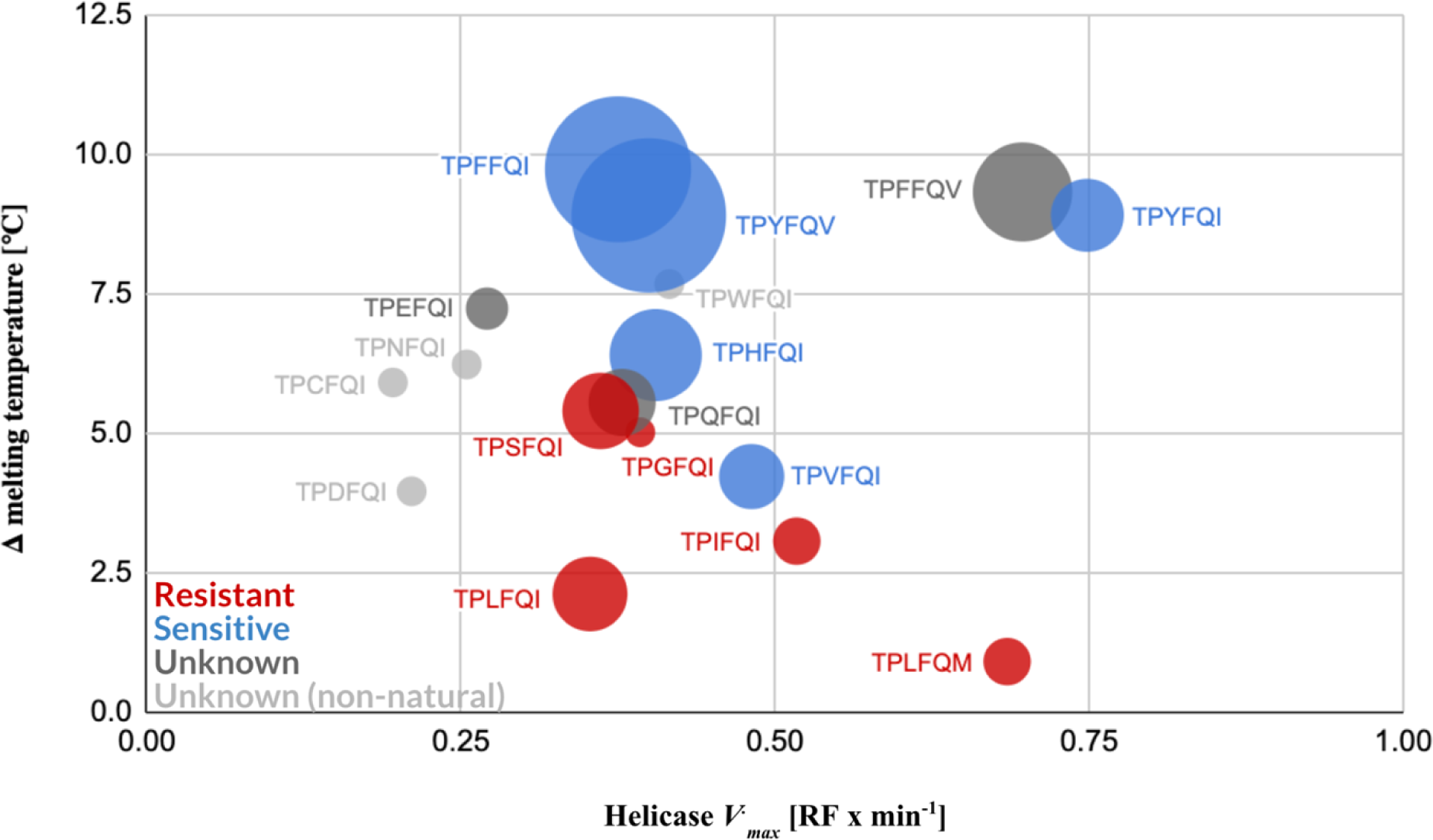
Shifts in thermal denaturation temperature of different eIF4A:RNA:silvestrol complexes are associated with eIF4A sensitivity to rocaglates. While relative eIF4A helicase activities of eIF4A mutant proteins expressing different rocaglate-binding aa patterns fall within a narrow range of *V_max_* values and do not correlate with rocaglate sensitivity, comparative thermal shift analysis of the mutant proteins showed a clear association between sensitivity to rocaglates and higher thermal denaturation differentials between the eIF4A:RNA and eIF4A:RNA:silvestrol complexes. The increased stability of the rocaglate-sensitive mutants is determined by π-π-stacking interactions elicited by the corresponding aa residues at position 163. Data points represent mean values of three technical replicates. Standard errors for the helicase activities are indicated in Fig. 4A and standard errors for the ***Δ*** melting temperature are listed in **Table S5**. The size of the circles denotes prevalence of the aa pattern among the eIF4As included in our survey.

As pointed out in the previous section, all the measurements presented here correspond to single or double aa variants generated on a human eIF4A1 protein scaffold. While such an analysis removes potential structural confounders, it also preempts an evaluation of the substitutions in positions 163 and 199 in their naturally evolved structural context, which could further contribute to the RNA and rocaglate binding characteristics of individual eIF4As. A comprehensive analysis of natural eIF4A variants was clearly out of the scope of this manuscript, but we have made efforts to address the dynamics of rocaglate binding to natural variants of eIF4A in a species-specific way. For instance, a preliminary thermal denaturation shift analysis of purified *Aedes aegypti* eIF4A1 (TPHFQV) with silvestrol has shown a ***Δ*** melting temperature of 8.45 °C, which is higher than the 6.41 ℃ observed for the TPHFQI human eIF4A1 variant (**Table S6**). An H163L mutant of the *Ae. aegypti* eIF4A further exhibited a reduction by 2.35 ℃ in the **Δ** melting temperature to 6.1 ℃, mirroring the trend in melting temperature changes observed for the same aa 163 mutations (F163H and F163L) in the human eIF4A1 protein scaffold (**Table S6**). While providing only a preliminary validation of our approach, these results also confirm the need to study the rocaglate sensitivity associated with specific substitutions of F163 in their naturally evolved context to obtain a comprehensive and accurate assessment of their effect on sensitivity to rocaglates and RNA helicase activity.

### Sensitivity to rocaglates can be predicted through combined binding energy inference and thermal denaturation shift measurement analysis of eIF4A:RNA:rocaglate complexes

To assess whether *in silico* inference of the binding energies and intermolecular contact levels of the eIF4A:RNA:rocaglate complex could help assess the potential sensitivity or resistance of a particular eIF4A to rocaglates, we performed a docking analysis of the eIF4A:RNA:rocaglate complex for all 35 aa patterns in the human eIF4A1 background in association with silvestrol, zotatifin, and CR-1-31-B [58, 41]. Both parameters were highly correlated with each other, and the lowest binding energies and highest intermolecular contacts also correlated with higher preponderance of the corresponding eIF4A variants in nature (**Figure S5**). Clade-specific analysis further showed evolutionary patterns whereby eIF4A variants that were shared with one or more other clades converged toward low binding energy/high intermolecular contact variants in all four groups—protists, fungi, plants, and animals (**Figure S6**).

Next, we explored whether a combination of *in silico* predicted parameters and *in vitro* measurements would help better differentiate rocaglate sensitive eIF4A variants from resistant ones. The combined analysis of thermal denaturation shift measurements and inferred binding energies provided the strongest separation between sensitive and resistant variants of eIF4A (**Figure 6A**). Similarly, we saw an association between intermolecular contacts and sensitivity to rocaglates when we analyzed this parameter in combination with thermal denaturation (**Figure 6B**).

**Figure 6:**
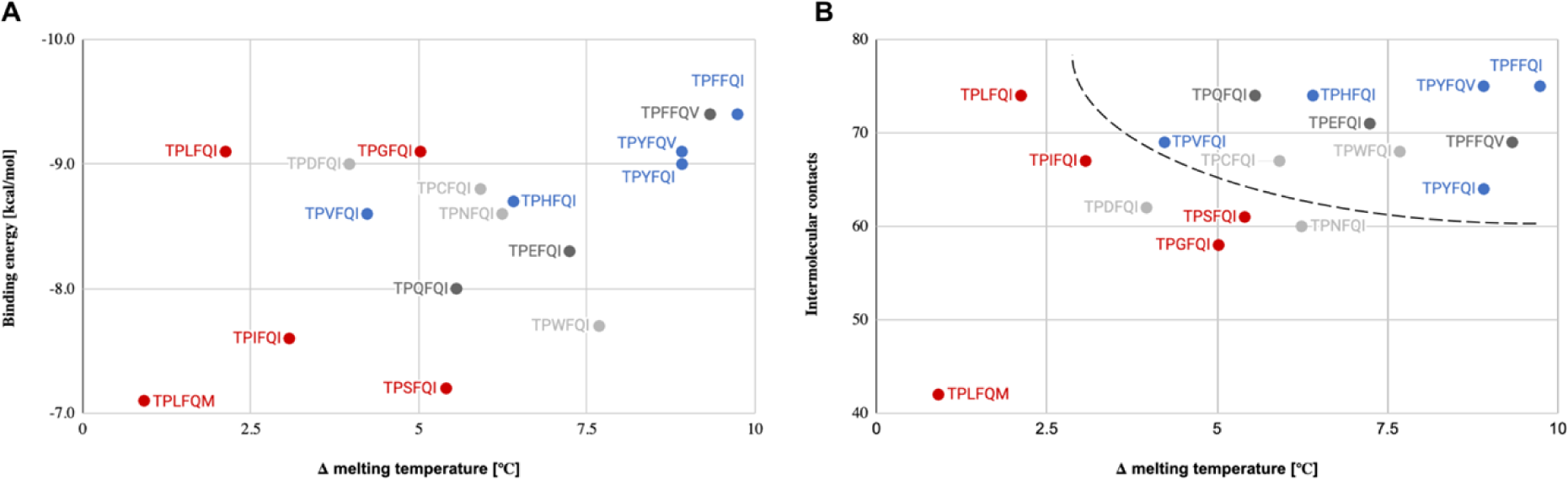
Combined binding energy or intermolecular contact inference with thermal denaturation shift measurement analysis of select eIF4A:RNA:silvestrol complexes reveals strong association with sensitivity to rocaglates. (**A**) Sensitivity to rocaglates (blue) was associated with high thermal denaturation shifts and low binding energies, while resistance (red) was associated primarily with low melting temperatures. Dark gray denotes natural eIF4A1 variants of untested resistance to rocaglates; light gray denotes non-natural eIF4A1 variants. (**B**) Sensitivity to rocaglates (blue) was associated with high thermal denaturation shifts and high intermolecular contacts, while resistance (red) was associated primarily with low melting temperatures. Dark gray denotes natural eIF4A variants of untested resistance to rocaglates; light gray denotes non-natural eIF4A variants; dashed line denotes approximate separation between rocaglate sensitive and resistant variants. Data points represent mean values of three technical replicates (***Δ*** melting temperature) and single values from optimized docking analysis (intermolecular contacts). Standard errors for the ***Δ*** melting temperature are listed in **Table S5**.

To further establish the strength of the associations observed with silvestrol, we conducted analogous analyses with the two synthetic rocaglates zotatifin and CR-1-31-B. A side-by-side comparison of the binding energies and thermal denaturation shifts determined for the three rocaglates revealed striking patterns of potential predictive power relative to rocaglate sensitivity (**Figure 7**). In particular, sensitivity to rocaglates was associated with shift patterns in melting temperature and in binding energies that were mirrored across the four variants we analyzed for which sensitivity has been shown *in vitro*. The pattern showed consistently higher melting temperature shifts and binding energies for silvestrol and practically overlapping values for these two parameters for zotatifin and CR-1-31-B. Another eIF4A variant, TPFFQV, which has not been shown *in vitro* yet to be sensitive to rocaglates exhibited an almost identical pattern to those of the known rocaglate-sensitive variants (**Figure 7**).

**Figure 7:**
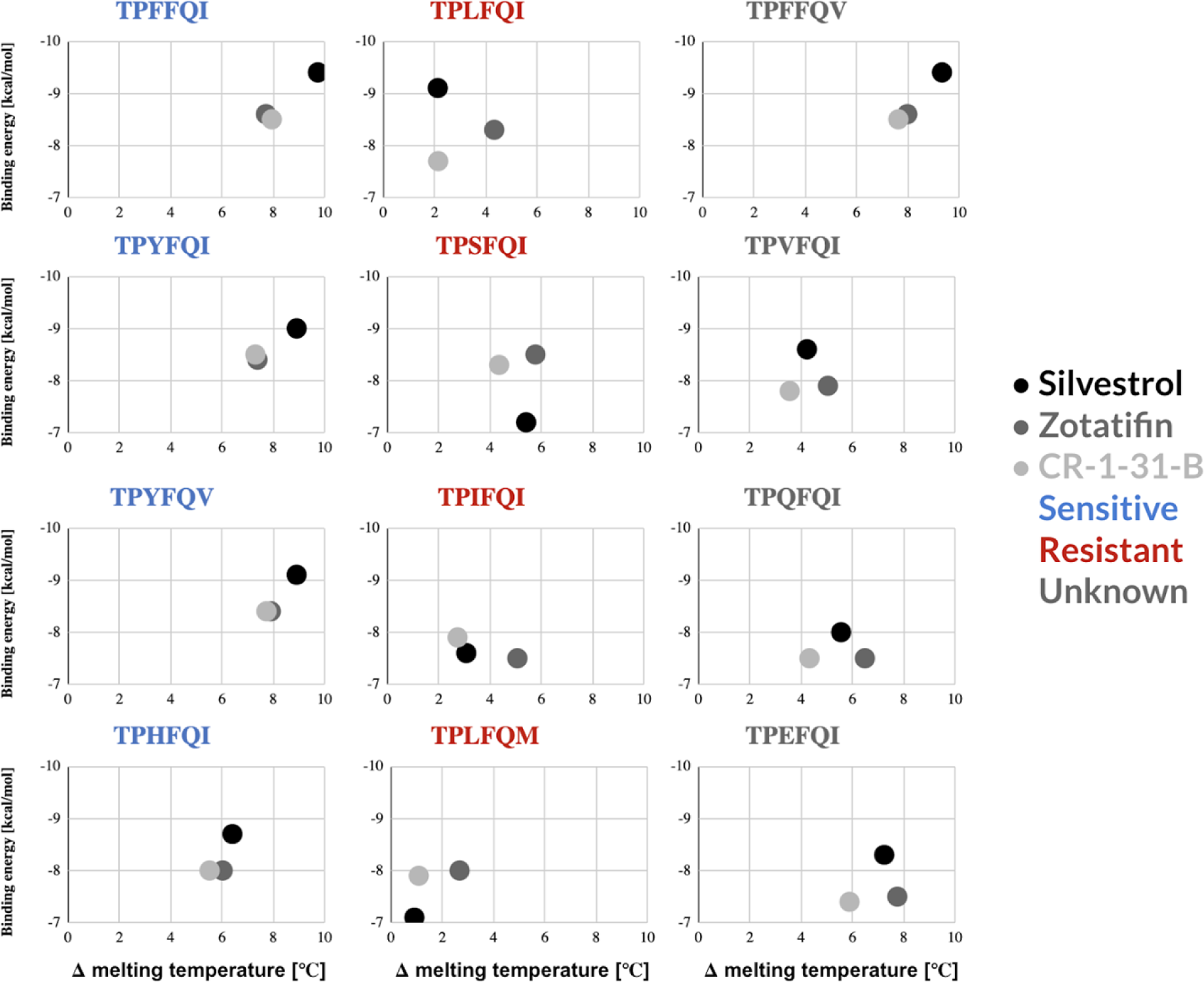
Side-by-side comparison of combined binding energy inferences and thermal denaturation shift measurements for silvestrol, zotatifin, CR-1-31-B. All eIF4A1 variants known to be sensitive to rocaglates (blue) exhibited remarkably similar patterns for all three rocaglates. By contrast, the four rocaglate-resistant variants analyzed here (red) showed widely disparate patterns. The four variants for which no experimental determination of sensitivity to rocaglates exists (dark grey), showed two distinct patterns—one that was analogous to that of the sensitive patterns analyzed here, and three that were analogous among them and different from all other sensitive or resistant patterns.

No comprehensive structure-activity analysis of the eIF4A:RNA complex with different rocaglates exists. To help explain the observed differences in melting temperatures, we performed initial structure-based computational modelling, which revealed that in addition to its interactions with select amino acids in the RNA-binding pocket, silvestrol may exhibit expanded intermolecular interactions with nearby Arg residues on the surface of eIF4A— Arg110, Arg282, and Arg311—via its unique 1,4-dioxane moiety (**Figure S7**). Each of these three Arg residues belongs to a different, structurally distinct and conserved motif of eIF4A: Arg110 is part of the PT**<u>R</u>**ELA sequence characteristic of motif Ia, Arg282 is part of the VIFCNT**<u>R</u>** sequence characteristic of motif IV, and Arg311 is part of the Qxx**<u>R</u>** motif. These three motifs fold into a highly conserved ‘Arg pocket’ that is adjacent to the RNA-binding pocket [45]. The Arg311 residue has also been shown to be involved in the formation of a critical salt bridge to the phosphate groups of RNA [41, 59].

Silvestrol’s interactions with this Arg pocket are likely to be mediated by its 1,4-dioxane moiety, a feature unique to silvestrol and its diastereomer episilvestrol (**Figure S1**). Interestingly, the crystal structure of the eIF4A:RNA:PatA complex has revealed that PatA, which almost perfectly mimics the interaction of rocaglates with the RNA-binding pocket, also establishes hydrophobic interactions via its trienyl amine moiety with Arg110 and Arg 282 as well as with Gly304 and Asp305, two highly conserved residues involved in RNA-binding and closely associated with the QxxR motif [55] (**Figure S7**).

Our initial mutagenesis studies of the Arg pocket further showed that single and triple substitutions of these three Arg residues to Ala result in a loss of ability of three rocaglates— silvestrol, RocA, or CR-1-31-B—to associate with eIF4A as a result of the reduced ability of RNA to form a complex with eIF4A in the first place (**Table S6**). Mutations in both position 110 and 282 result in reduced melting temperature shifts—R282A reduces the temperature shift in half while R110A results in a neutralization of the temperature shift. Given that no addition of RNA to the thermal denaturation assays results in a consistent destabilization of the eIF4A association by about -3.32 ℃ (SD ±0.53) (**Table S6**), the temperature shifts we observed for the R110A and the R282A suggest a reduced ability of the mutated eIF4A proteins to bind RNA. By contrast, the R311A and the triple mutants exhibited equivalent negative temperature shifts to those observed in all control assays without RNA addition, indicating an inability of the mutated eIF4A proteins to bind RNA (**Figure S8**).

In contrast, the picture for the rocaglate-resistant eIF4A variants showed no clear relative binding patterns for the three rocaglates we tested (**Figure 7**). Three of the four variants exhibited different substitutions at position 163—Leu, Ile, and His—against a constant human eIF4A1 background, and one variant—TPLFQM—contained both a Leu substitution at position 163 and a Met residue at position 199. The extreme values of the experimentally and *in silico* determined values for the TPLFQM variant, including its optimal RNA helicase *V_max_*,, point to this variant as an evolutionarily favored variant in the context of *Aglaia* sp..

Besides the TPFFQV variant, the three other natural eIF4A variants of unknown sensitivity we analyzed—TPVFQI, TPQFQI, and TPEFQI—exhibited analogous patterns among them, with lower binding energies for silvestrol than for zotatifin and CR-1-31-B, and consistently higher melting temperatures for zotatifin than for silvestrol or CR-1-31-B (**Figure 7**). Based on the binding energies and intermolecular contacts inferred from our docking analysis, the Val residue at position 163 could mediate rocaglate-triggered clamping via hydrophobic interactions between phenyl ring C of the rocaglates and the aliphatic chain of Val, making this pattern sensitive to rocaglates (**Figure 8**). Interestingly, the carbon backbone of Val is shorter than those of Leu and Ile, both of which have been shown to prevent the formation of π-π-stacking or hydrophobic interactions and did not allow for silvestrol to interact with eIF4A in our docking analysis (**Figure 8**). Our docking analysis predicted that a Val residue in position 163 might favor the establishment of stable hydrophobic interactions with rocaglates, but this would have to be tested.

**Figure 8:**
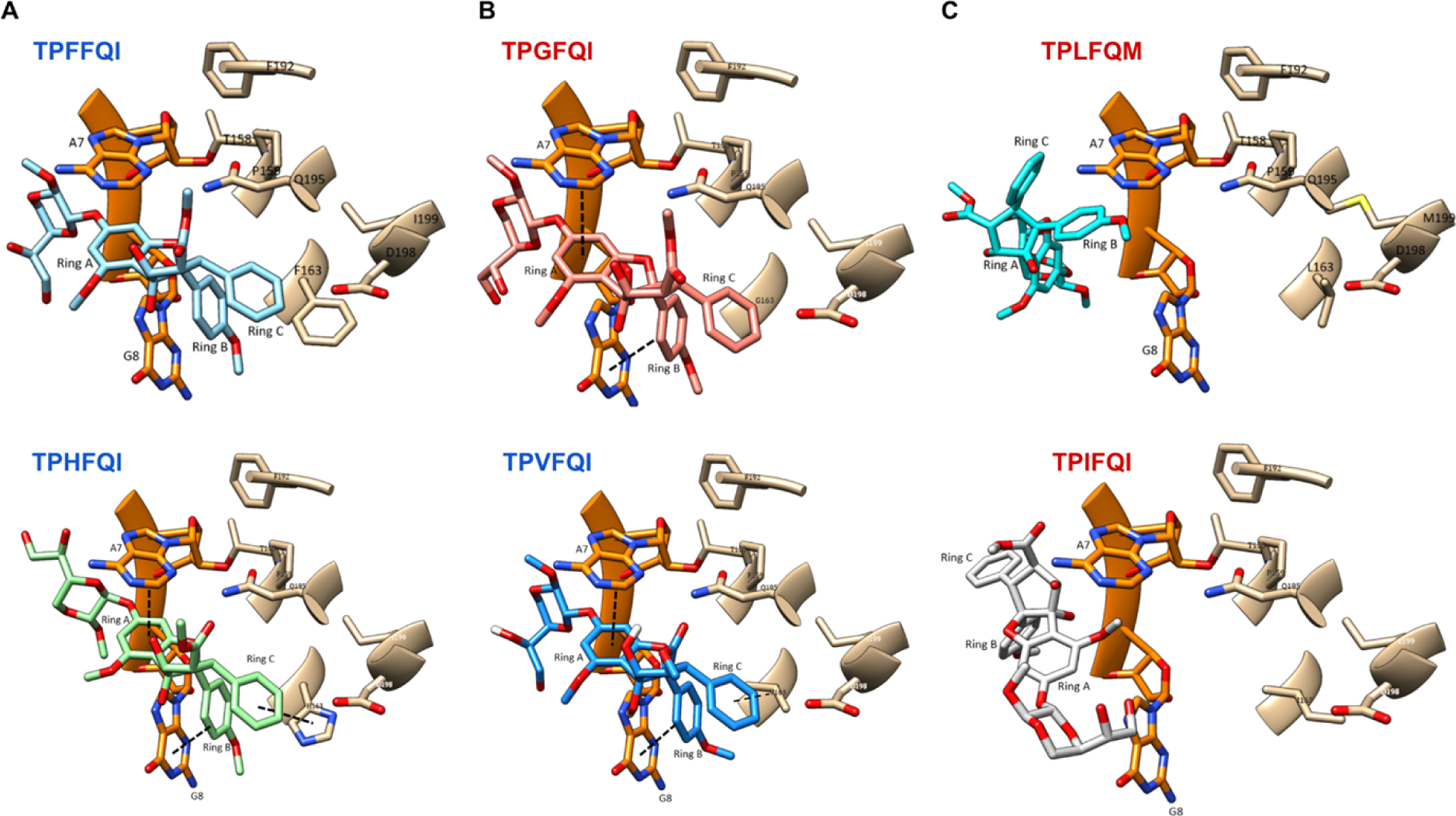
Molecular docking of silvestrol to select eIF4A variants. (**A**) Aromatic or aromatic-like aa residues at position 163 such as Phe or His, respectively, allow for the establishment of stable π-π-stacking interactions with rocaglates. (**B**) Substitutions with short chain aa residues such as Val allow for hydrophobic interactions with rocaglates that can to some extent compensate for the lack of π-π-stacking interactions. (**C**) Long aliphatic side chains such as Leu or Ile sterically preempt the formation of stable hydrophobic interactions, rendering the corresponding eIF4As resistant to rocaglate mediated clamping of the eIF4A:RNA complex.

The docking analysis of both the TPQFQI and the TPEFQI mutants predicted the formation of stable hydrophobic interactions with rocaglates in analogous conformations to the π-π-stacking observed with the two aromatic residues Phe and Tyr or the aromatic-like basic aa His in position 163 (**Figure 8**). Given the radically different nature of Gln, an amine residue, and Glu, an acidic residue, it was interesting to observe their similarity in binding energies and thermal denaturation shift patterns vis-á-vis silvestrol, zotatifin, and CR-1-31-B. This similarity, which implies the establishment of stable hydrophobic interactions and thus corroborates the docking analysis, is most likely due to the steric conformation determined by the long carbon backbones of Gln and Glu.

### *In vitro* assays confirm predicted sensitivities to silvestrol

To confirm predictions of sensitivity or resistance to rocaglates based on the outcomes of our molecular docking and temperature shift analyses, we performed *in vitro* silvestrol sensitivity assays with four species expressing eIF4As with previously untested aa patterns, four species with previously tested aa patterns but belonging to new genera, and one species belonging to a genus that had been previously characterized as rocaglate sensitive but had not been tested with silvestrol (**Table 1**).

**Table 1:** List of organisms used for *in vitro* testing of silvestrol sensitivity. (* motif inferred from 100% consensus among 32 fruit flies across five genera)

| Species | Predicted sensitivity | Rocaglate-binding aa pattern | Measured sensitivity [S – sensitive; R – resistant] | Assay |
| --- | --- | --- | --- | --- |
| *Schistosoma mansoni* | Sensitive | TPFFQI | S | Motility and egg production assays |
| *Anastrepha suspensa* * [AsE01] |  | TPYFQV | S | Cell proliferation assay |
| *Drosophila melanogaster* [S2] |  |  | S | Cell proliferation assay |
| *Plasmodium falciparum* [3D7] |  |  | S | Viability assay in erythrocytes |
| *Aedes aegypti* [Aag2] |  | TPHFQV | S | Cell proliferation assay |
| *Toxoplasma gondii* |  |  | S | Proliferation assay in MARC145 monkey kidney cells |
| *Trypanosoma brucei brucei* [Lister 427] |  | TPVFQI | S | Viability assay |
| *Leishmania amazonensis* [LV78] | Resistant | TPSFQI | R | β-lactamase assay in J774 macrophage cells |
| *Caenorhabditis elegans* |  | TPGFQV | R | Developmental and lifespan assays |

We selected two pathogens—*Toxoplasma gondii* and *Trypanosoma brucei brucei*—the pathogen vector *Aedes aegypti,* and the non-pathogenic nematode *Caenorhabditis elegans* to test whether our predictions on eIF4A sensitivity to rocaglates based on the docking and thermal shift analyses could be confirmed. *Ae. aegypti*, *T. gondii* and *T. brucei brucei* were predicted to be sensitive to silvestrol based on their aa substitutions at position 163, Phe to His (*Ae. aegypti*, *T. gondii*) and Phe to Val (*T. brucei brucei*), and *C. elegans* was predicted to be resistant to silvestrol based on its F163G substitution. Using model-specific viability, developmental or lifespan assays, we were able to confirm all three predictions (**Figures S9 & S10**).

We complemented the series by testing another important pathogen, *Schistosoma mansoni*, and the non-pathogenic fruit flies *Anastrepha suspensa* and *Drosophila melanogaster*. The eIF4A sequences of *S. mansoni* and *D. melanogaster* contained aa patterns TPFFQI and TPYFQV, respectively, which had been previously shown to be sensitive to rocaglates. The eIF4A sequence of *A. suspensa* was not available at the time of this writing, but the eIF4A mRNA sequence of the closely related *A. fraterculus* was available (TPYFQV) [60] and, given the 100% consensus of this motif across five genera and 32 species of fruit flies (**Table S2**), we assumed the presence of the same motif in *A. suspensa*. With these assays we were able to confirm the applicability of sensitivity results from one species to unrelated genera. In addition to *T. brucei*, we also tested two other protozoan pathogens—*Leishmania amazonensis* and *Plasmodium falciparum*—to confirm that aa patterns associated with silvestrol sensitivity in one species would confer sensitivity to different species within the same genus containing the same aa pattern. We also observed that two protozoans within the same family—*T. brucei* and *L. amazonensis*—exhibited rocaglate sensitivity and resistance, respectively, in concordance with their corresponding eIF4A aa patterns (**Figures S10 & S11**; **Table 1**).

All assays confirmed the predictions, expanding the number of potential pathogens that can now be targeted with rocaglates from the 50% we initially estimated based on known sensitivity reports to 60%, including *Cystoisospora suis*, the most common pathogen affecting suckling and weaned piglets, and the major wildlife pathogen *Aphanomyces astaci*, a globally distributed protist responsible for freshwater crayfish plague (**Table S2**). The picture around resistance to rocaglates also becomes clearer, raising the proportion of potentially resistant pathogens from 13% to 19%, and now including agriculturally important plant and animal pathogens such as *Phytophthora nicotianae* and *Aphanomyces invadans* (**Table S2**).

The combined binding energy and thermal denaturation shift analysis showed analogous patterns for three variants of unknown sensitivity to rocaglates: TPVFQI, TPEFQI, and TPQFQI (**Figure 7**). The *in vitro* results revealing the sensitivity of TPVFQI to silvestrol suggests that organisms containing the analogous aa combination TPEFQI and TPQFQI could also be sensitive to rocaglates. This would add further organisms to the list of pathogens and other harmful organisms potentially targetable with rocaglates such as the fungus *Encephalitozoon cuniculi* (TPEFQI), a causative organism of neurologic and renal disease in rabbits and sometimes immunocompromised humans, and important plant pests and plant pathogens such as *Bactrocera latifrons*, *Aspergillus fischeri*, *Marssonina brunnea*, and *Zopfia rhizophila*, all of which express the TPQFQI variant of eIF4A (**Table S2**).

## Discussion

In the ongoing search for novel ways to fight non-viral and non-bacterial pathogens, targeting translation to inhibit growth and overall viability has emerged as an attractive strategy. Here, we focused on the potential of rocaglates, a group of plant-derived compounds that target the DEAD-box RNA helicase eIF4A, a key component of the eukaryotic translation initiation complex eIF4F. Our analysis uncovered a large proportion of pathogens potentially targetable with rocaglates, providing actionable information for prioritization and development strategies.

The eIF4A helicase is one of the most highly conserved proteins in eukaryotes, possibly as a result of its critical role in translation as an essential enzyme for processing mRNAs containing stable RNA secondary structures in their 5’UTRs. Our analysis showed in more detail how four out of the six residues involved in the interaction of eIF4A with rocaglates are indeed strictly conserved across clades in the tree of life, likely a result of their structural role within the RNA-binding pocket of eIF4A. Conversely, the remaining two aa residues in positions 163 and 199 showed startling substitution tolerance patterns that among others determine sensitivity to rocaglates. Position 163 in particular exhibited a remarkable tolerance for aa substitutions that cover a broad range of aliphatic, aromatic, basic, acidic and other aa. The distinct distributions of aa substitution frequencies at this position further indicate a level of ‘evolutionary preference’ across eukaryotic groupings suggestive of a certain directionality in evolution towards a preponderance of Phe and Tyr. Position 199 showed a more constrained tolerance, with a bimodal distribution of Ile and Val throughout eukaryotic groupings, and a unique Met substitution in *Aglaia* species that produce silvestrol and/or episilvestrol.

RNA helicase activity of select variants expressed in a human eIF4A1 background showed remarkable consistency in relative *V_max_*, supporting the notion that substitutions at the six rocaglate-binding aa residues are under evolutionary pressure based on their effect on the RNA helicase activity of eIF4A rather than its sensitivity to rocaglates. The comparable *V_max_* of the two eIF4A isoforms found in *Aglaia* confirms this notion.

Our analysis reveals that, *a priori*, a high proportion of known pathogens should be targetable with rocaglates, opening up new space for targeted development of anti-pathogen agents. The expanded evolutionary picture of natural rocaglate resistance, together with some novel mechanistic insights into the molecular interactions between diverse rocaglates and different eIF4A variants, further provide important landmarks for developing rocaglate derivatives or novel compounds as well as sound implementation strategies that minimize the risk for emergence of *de novo* resistance.

### Enlisting rocaglates to fight eukaryotic pathogens

Our analysis revealed the potential for using rocaglates and derived molecules to combat a large proportion of known human, plant, and animal pathogens. The unusually well-defined determinants of eIF4A sensitivity to rocaglates—the six aa residues analyzed in our study— provide a precise tool for predicting potential effectiveness of novel compounds against a pathogen of interest. Our mutant analysis has further expanded the spectrum of potential targets by providing the first *in vitro* confirmation for three new variants: TPVFQI, and TPHFQV, which render pathogens sensitive, and TPGFQV, which provides resistance to rocaglates.

The TPVFQI variant, characteristic of *Trypanosoma* sp., opens the possibility of addressing important human diseases such as sleeping sickness, caused by *T. brucei*, and Chagas disease, caused by *T. cruzi*. The TPHFQV variant is found in a number of important agricultural pathogens including *Cystoisospora suis*, a parasite that causes diarrhea with a high mortality rate in piglets, *Marssonina coronariae*, an apple blotch–causing fungus that results in severe premature defoliation, and, most importantly, in *Aedes sp.*, a group of widely distributed and high-impact disease vectors that includes the yellow fever mosquito, *Ae. aegypti*, and the Asian tiger mosquito, *Ae. albopictus*. The rocaglate-resistant variant TPGFQV, which with the exception of *Caenorhabditis elegans*, a nematode, is only found in protists, rules out the use of rocaglates or its derivatives for a small but diverse number of pathogens that includes the plant pathogen *Phytophthora* sp., the causative agents of several plant blights including potato and tomato blights, *Aphanomyces* sp., a causative organism of epizootic ulcerative syndromes on many species of fish and shellfish, and *Saprolegnia sp.*, a group of pathogens that affects fish and amphibians worldwide. The analogous variant TPGFQI had previously been shown to be resistant and has so far only been identified in the *Aglaia* sp. fungal parasite *Ophiocordyceps* sp..

Mechanistically, eIF4A sensitivity to rocaglates is determined by the formation of stable π-π-stacking interactions between the rocaglates and the RNA-binding pocket of eIF4A through interactions with the rocaglate-interacting residues analyzed here. Our sensitivity findings, as well as those from previously published sensitivity studies, could be predicted by our *in silico* modeling of binding energies and intermolecular contact levels of the ternary eIF4A:RNA:rocaglate complex and by our *in vitro* analysis of the melting temperatures of the different eIF4A:RNA complexes.

However, while our comprehensive analysis of all known eIF4A sequences from pathogens provides a first glimpse at the potential for enlisting rocaglates to fight a range of human, animal and plant pathogens, several important questions remain to be answered.

First, the diversity of eIF4A variants we have revealed, while informative, is likely limited in terms of the true diversity of variants existing in nature. We analyzed a curated list of 365 eIF4A sequences, many of which belonged to related species within the same genus. The analysis revealed 35 variants with respect to the six rocaglate-interacting aa residues we focused on here. Clearly, this only amounts to a drop in the bucket in terms of potential diversity in nature, including among hitherto not-sequenced pathogens. Nonetheless, our study provides a first insight into the potential evolutionary constraints that have determined resistance to rocaglates in the past and could drive it in the future under increased exposure to the compounds.

Secondly, and while eIF4A is one of the most conserved proteins known to us, it is also true that even small changes in other parts of the helicase could affect the tertiary structure of the RNA-binding pocket and thus its ability to interact with rocaglates in just the right orientation to allow for the formation of the necessary π-π-stacking interactions. Our mutant work has shown the effect of single aa substitutions on intermolecular dynamics, but these were all determined against a fixed human background. How those variants perform in their natural protein scaffolds cannot be predicted with 100% confidence based on our studies alone and would necessitate *in vitro* analysis of the natural proteins.

And thirdly, the interaction of different rocaglates with eIF4A varies in subtle ways that are not yet completely understood. Our analysis revealed how these interactions can sometimes be predicted from the rocaglate structures alone—silvestrol exhibits overall higher melting temperatures than zotatifin or CR-1-31-B probably due to its 1,4-dioxane moiety, which could provide an expanded interface for contacts with nearby arginine residues of eIF4A. In other instances, however, and depending on the motif they are interacting with, the binding dynamics paint a more complex picture—the TPIFQI variant elicits analogous interactions with silvestrol and CR-1-31-B that are distinct from those with zotatifin and can thus not be solely explained by the presence of silvestrol’s 1,4-dioxane moiety. Understanding these natural interaction dynamics will provide further insight into how to design rocaglates of improved efficacy and specificity.

### Does natural resistance to rocaglates provide a fitness advantage?

Our global eIF4A variant analysis in the context of the eukaryotic tree of life—a widely accepted proxy for a rough evolutionary progression of the main eukaryotic groups of organisms—revealed a random distribution of resistance variants across protists, fungi, plants and animals. Importantly, known rocaglate biosynthesis is limited to one genus of plants, *Aglaia*, which would have emerged at a much later stage than most of the resistant variants we detected. While it is impossible to categorically determine the trigger for the emergence of rocaglate-resistant eIF4A variants, and given that a potential functional role of this alkaloid in *Aglaia* sp. is not known, two scenarios can be explored with our results.

In a first scenario, resistance would have emerged as a direct consequence of exposure to rocaglates following the appearance of rocaglate biosynthesis in *Aglaia*. A close analysis of the list of resistant organisms reveals a number of pathogenic and non-pathogenic organisms that exhibit geographical distributions supportive of such a scenario. *Aglaia* has a mostly East and Southeast Asian distribution, which overlaps with the geographical distributions of resistant organisms such as the river fluke *Clonorchis sinensis*, oriental fruit flies *Bactrocera sp.*, and the flatworm *Dugesia japonica*, all three known regional pests or pathogens, or the Pacific oyster *Crassostrea gigas*, a non-pathogen. More immediately, the recent description of the *Aglaia* sp. parasitic fungus *Ophiocordyceps* sp. BRM1 also supports the potential emergence of resistance following direct exposure to rocaglates. However, there are many other rocaglate-resistant organisms that have non-overlapping distributions with that of *Aglaia*, or global distributions, making it unlikely that rocaglates provided an evolutionary pressure. Tracing the fossil record of *Aglaia* and the organisms associated with it over evolutionary time would be one way of addressing this question, but was out of the scope of our work.

In a second scenario, rocaglate resistance would have been an inconsequential byproduct of natural eIF4A variation. This scenario would be supported by *Aglaia sp.* being the only organisms in which both eIF4A isoforms contain rocaglate-binding amino acid variants resistant to rocaglates, TPLFQM and TPLFQI, and by the fact that the relative *V_max_* of the different natural eIF4A variants we analyzed was remarkably constant. The plants’ ability to produce rocaglates and the inherent tolerance for substitutions at positions 163 and 199, could have led to the establishment of two rocaglate-resistant eIF4A isoforms because this would provide a clear fitness advantage to the plants. Previous work has shown that while both eIF4A isoforms exhibit equivalent biochemical activities, eIF4A1 is essential for cell survival, while eIF4A2 is not [12].

The emergence of rocaglate biosynthesis in *Aglaia sp.* would have thus been accompanied by the establishment of the necessary ‘resistance’ mutations in eIF4A to protect the plants from self-poisoning. The rocaglate-resistant aa pattern of the main eIF4A isoform in *Aglaia*, TPLFQM, contains the most common ‘resistant’ substitution at position 163, Leu, and a Met substitution at position 199, a substitution we have shown increases both the *V_max_* of the helicase as well as substantially decreases the thermal denaturation temperature of the eIF4A:RNA:rocaglate complex.

None of these scenarios can be ruled out because both the aa substitution tolerance at positions 163 and 199 could be driving the random appearance of ‘resistant’ variants and the physical proximity and exposure to rocaglates could be triggering the retaining of ‘resistant’ variants in select organisms simultaneously. However, understanding both processes will be critical for the successful implementation of any rocaglate-based anti-pathogen measures.

### Thoughts on the emergence of *de novo* resistance to rocaglates in pathogens

Our analysis has laid out the landscape of potential pathogen targets for rocaglates based on the limited spectrum of eIF4A sequences that have been obtained to date through mostly whole genome sequencing of organisms of interest for human, animal and plant health. While clearly incomplete in terms of representing the total diversity of eIF4A sequences in nature, our analysis revealed very clear conservation patterns in four out of the six positions relevant to rocaglate binding—158, 159, 192, and 195—and equally significant substitution tolerance patterns in the other two positions—163 and 199. With our analysis of natural and non-natural mutants, we aimed to further complete the picture of potential variants and their rocaglate-binding dynamics. We believe this closer-to-complete picture of the sequence diversity already established in nature provides a glimpse into the potential for the development of rocaglate resistance and the need for proper management of any potential implementation of rocaglates as anti-pathogen agents.

Our experience over the last less than hundred years deploying antimicrobials to fight infection has taught us that, when faced with a new challenge, nature will always adapt through natural evolution determined and driven by the survival of the fittest, in this case the variant that exhibits a fitness advantage [61]. In most cases, this adaptation has resulted in the development or acquisition of resistance mechanisms to antimicrobials, whether through lateral transfer of genes from other microorganisms, *de novo* mutations, or repurposing of existing mechanisms for detoxification.

The emergence of resistance to pesticides has also been well studied. Unlike with microbes, where short generation times, *de novo* mutations, and lateral gene transfer play major roles in the emergence of resistance traits, emergence of resistance in more complex organisms such as fungi and insects has been tied to factors including the number of point mutations needed for resistance, the pre-existence of resistance alleles in a population, and the fitness of the mutated resistant variants in the absence of the corresponding selection pressure [62, 63].

Our analysis points to the possibility that resistance of eIF4A to rocaglates could be an accidental trait arising from the aa substitution tolerability at position 163 of eIF4A. Single point mutations leading to aa changes at this position can turn a rocaglate-sensitive organism into a resistant one. While this could be categorized as a single aa mutation challenge, our analysis has confirmed previous work that suggested the steric constraints of the RNA-binding pocket of eIF4A determined the high level of sequence conservation of this enzyme. Rocaglate- driven clamping is mainly determined by the aa residue at position 163, however, this has to happen within the constraints of four other residues having to remain unchanged—158, 159, 192, and 195—and one being tolerant to minimal substitution—199. In addition, clamping is also highly sensitive to nucleotide composition, requiring the presence of two adjacent purine RNA bases to stabilize the eIF4A1:RNA complex, which in turn constrains the tolerability for substitutions at the relevant RNA interacting aa residues of eIF4A [41]. The combination of these factors thus makes the emergence of resistance a more complex albeit overall still low barrier evolutionary event.

Our analysis has further shown that nature has already evolved a sizable number of rocaglate-resistant eIF4A alleles, with some of them already co-existing with sensitive isoforms in some organisms. This would seem to *a priori* pose a major barrier to the implementation of rocaglate-based anti-pathogen strategies. However, the availability of this extensive catalog of information about sequence, structural and physicochemical factors that characterize ‘resistant’ alleles could also be used to inform the development of novel compounds, the deployment of carefully managed control programs, and the monitoring of the emergence of potential *de novo* resistance in managed populations.

A key determinant for the *de novo* emergence of resistance to rocaglates is the potential fitness advantage associated with the resistant versus the sensitive variants. Within the genus *Aglaia*, this tolerability was an asset when it came to adapt to the emergence of its own capability to synthesize rocaglates, as proven by both isoforms of eIF4A in these plants being resistant to rocaglates. How this process could play out in other species only transiently exposed to rocaglates is not known. ‘Fitness rescue’ in species containing a resistant and a sensitive isoform has so far only been shown in cases where isoform 1 is the resistant isoform but not when it is isoform 2 that contains the resistant allele. This is the case for most of the pathogens we have analyzed so far. And while we have no way of knowing how fast resistant mutations have been acquired over evolutionary timescales, our analysis has revealed that, at least in theory, resistant mutations could arise quickly through point mutation events at the codon wobble position as exemplified by the switch from Phe to Leu.

As shown by our eIF4A helicase activity analysis, all natural variants of eIF4A exhibit relative *V_max_* within a narrow range that is most likely essential for the fitness of the organism. This applies to both resistant and sensitive variants of eIF4A, suggesting that fitness would not be a deciding factor for the permanent establishment of a resistant allele because there would be no detrimental fitness effects in the absence of rocaglates.

The combined evaluation of these three aspects—the minimal number of aa substitutions needed for resistance, the pre-existence of resistance alleles in the population, and the fitness of the mutated resistant variants in the absence of rocaglates—paints a complex picture of opportunity for the anti-pathogen application of rocaglates. While the aa substitution tolerance levels at position 163, the wide distribution of natural resistance alleles, and the *a priori* neutral fitness effects of resistance all seem to compromise the potential of harnessing rocaglates for managing pathogens, this knowledge, which was often non-existent before the implementation of other anti-pathogen compounds, could provide the basis for more robust and sound implementation strategies. Measures such as punctual, high concentration deployments of rocaglates accompanied by comprehensive monitoring programs could be one approach to minimizing the risk for triggering the emergence of resistance. Taking advantage of the favorable therapeutic windows of rocaglates in humans and animals compared to the sensitivity in fungi and protists could be pivotal in terms of developing safe interventions against select rocaglate-sensitive parasites. Harnessing the comprehensive catalog of natural eIF4A variants uncovered in our study could further help advance the search for novel synthetic rocaglates or other small molecule inhibitors of enhanced specificity and efficacy.

Overall, our study has provided a first comprehensive account of the natural diversity of rocaglate-resistant eIF4As and their distribution among pathogens, further physicochemical and rocaglate-binding characterization of select and highly represented variants, *in vitro* validation of several of our rocaglate-sensitivity predictions in pathogens, and further insights into the biological and evolutionary determinants of eIF4A-based rocaglate sensitivity.

## Methods

### eIF4A Sequence Analysis

Sequence analysis was performed using the full eIF4A protein sequences retrieved from <u>Genbank</u> and cross-validating them on <u>UniProt</u> to determine specific isoforms. Only sequences corresponding to the eIF4A1 and eIF4A2 isoforms were used for the analysis. Next, we extracted the six amino acid motifs associated with rocaglate binding for each of the protein sequences (human positions 158, 159, 163, 192, 195, 199) [41, 42]. Variant motif distributions were determined within each eukaryotic grouping and mapped onto the latest version of the eukaryotic tree of life [54]. Sequence logos illustrating amino acid tolerances for the six amino acids analyzed were rendered using Seq2Logo - 2.0 [53].

### eIF4A Variant Cloning, Overexpression and Purification

Select eIF4A variants with single and double substitutions at positions 163 and/or 199 were generated using PCR-based site-directed mutagenesis of a plasmid encoding human eIF4A1 (pET-28a(+)_eIF4A1(19-406) containing an *N*-terminal His-Tag and a thrombin cleavage site; variant-specific primers listed in **Table S4**). Double mutants were generated by first generating the mutations at position 163 followed by the mutations at position 199. Ligation products were transformed into *E. coli* DH5α cells and the plasmids were sequenced to confirm the corresponding substitutions. Following sequence confirmation, competent *E. coli* BL21 (DE3) cells were transformed with the plasmids and grown in lysogeny broth medium at 37 °C to OD_600_∼0.5. After the addition of 0.5 mM IPTG, the cells were grown at 15 °C for 16 h. The collected cells were lysed by sonication in a 20 mM HEPES-KOH buffer (pH 7.5, 300 mM KCl, 20 mM imidazole, 5mM ß-mercaptoethanol, 0.1 mM EDTA, 10 % (v/v) glycerol) with 1x cOmplete™, Mini, EDTA-free Protease-Inhibitor-Cocktail (Roche). The lysate was fractionated on a HisTrap^TM^ HP 1 mL column (GE Healthcare) using a linear gradient from sonication buffer to elution buffer (sonication buffer with 250 mM imidazole). The peaked fractions were collected, buffer-exchanged to a 20 mM HEPES-KOH buffer (pH 7.5, 300 mM KCl, 5 mM MgCl_2_, 0.1 mM EDTA, 1 mM DTT and 10 % (v/v) glycerol), flash-frozen in liquid nitrogen and stored at –80 °C.

### Helicase Assay

Helicase activities of the eIF4A variants were determined using a fluorescence-based assay. The capacity to unwind dsRNA substrates was measured using two labelled RNA substrates: a 10mer modified with Cyanine 3 (10mer-Cy3; 5’-[CY3]GCUUUCCGGU-3’), and a 16mer modified with Black Hole Quencher 2 (16mer-BHQ2; 5’-ACUAGCACCGGAAAGC[BHQ2]- 3’). An unlabeled competitor (10mer-competitor; 5’-GCUUUCCGGU-3’) was used to capture released quencher RNA. A single-stranded Cy3 RNA substrate (ssRNA) was used to determine the maximum fluorescence signal of the reaction. Equimolar amounts of 10mer-Cy3 and 16mer-BHQ2 were annealed at 80 °C for 5 minutes and incubated at room temperature for 1 hour followed by incubation on ice for 10 minutes in a 25 mM HEPES (pH 7.4 (KOH) in ddH_2_O). Competitor RNA was added in 1:10 (v/v) excess to the labelled RNA substrates and the reaction was again incubated on ice for 10 minutes prior to adding it to the helicase reaction mix. eIF4A (25 µM final concentration) was added to the reaction and fluorescence was measured using a Safire 2 microplate reader (Tecan).

### Thermal Shift Assay

Thermal shift assays were performed by incubating 5 μM of recombinant human eIF4AI (19- 406) with 50 μM of a polypurine RNA (AG)_5_ (Biomers, Ulm, Germany), 1 mM AMP-PNP (Roche, Basel, Switzerland), 100 μM of rocaglate (silvestrol, RocA, zotatifin, or CR-1-31-B) and 75 μM SYPRO Orange (S6650, Invitrogen, Carlsbad, CA, USA) in a 20 mM HEPES-KOH buffer (pH 7.5, 100 mM KCl, 5 mM MgCl_2_, 1 mM DTT, 0.1 mM EDTA, and 10% (v/v) glycerol) for 10 min at RT. The melt curves were measured between 10 ℃ and 95 ℃ at a 1.6 ℃/min ramp rate using the QuantStudio3^TM^ Real-Time PCR system (Applied Biosystems, Waltham, MA, USA) in a MicroAmp^TM^ Fast Optical 96-well plate (Applied Biosystems, Waltham, MA, USA).

### Docking Analysis

Molecular docking was performed using AutoDock, v4.2 [64]. The proteins were processed by adding all hydrogen atoms and merging non-polar hydrogen atoms using AutoDock Tools 1.5.7. Charges were assigned using the Gasteiger method with fixed torsions for the ligand. We set a 60 × 60 × 60, 3.75 Å grid box around the active sites with x, y and z-dimensions of 46.355, 9.919, 47.473, respectively. The rigid grid box was set using AutoGrid 4, followed by AutoDock with the Lamarckian genetic algorithm to obtain the best docking poses [65]. Dockings were performed in duplicate and the average binding energy reported. Select poses representing optimal binding affinities were visualized using UCSF Chimera (University of California).

### Cell-Based *In Vitro* Studies

#### Aedes aegypti, Anastrepha suspensa, <u>*and*</u> Drosophila melanogaster

Cell proliferation assays were performed with insect cell lines (Aag2 [*A. aegypti*], AsE01 [*A. suspensa*], and S2 [*D. melanogaster*]) using the WST-1 assay (Sigma-Aldrich). Cells (Aag2: 1.2 x 10^5^ cells/100 μL, AsE01: 1 x 10^5^ cells/100 μL, S2: 6 x 10^4^ cells/100 μL) were incubated in the presence of silvestrol (0 nM to 1.6 µM). Following a 24 hour incubation with silvestrol, we added WST-1 reagent as specified by the manufacturer and waited an additional three hours before determining cell mortality by measuring absorbance at 440 nm (reference wave length: 600 nm). CC_50_ values were determined for each set of biological replicates measured (GraphPad Prism V9).

### Whole Organism-Based *In Vitro* Studies

#### Toxoplasma gondii

The effect of silvestrol treatment on *Toxoplasma gondii* replication in MARC-145 cells was determined at 48 h post infection (h p. i.). Cell viability was controlled after 48 h of treatment with up to 100 nM silvestrol via XTT assays (solvent: DMSO (1:500); positive control: Triton X-100 treatment (1:200); negative control: plain medium). At 48 h p. i., the number of *T. gondii* tachyzoites released from infected host cells into the cell supernatant was determined. Assays were performed in triplicate.

#### Trypanosoma brucei brucei

Viability assays of *T. brucei brucei* (non-recombinant 427 strain) were performed for 48 h using HMI-9 medium (modified DMEM (IMDM; Cell Gro); 10% FBS; 10%, Serum plus (SAFC); 0.05 mM Bathocuproinesulfonate; 1.5 mM L-cysteine; 1 mM hypoxanthine; 0.2 mM *β*-mercaptoethanol; 0.16 mM thymidine; 1mM pyruvate). After a 48 h incubation, cells were labeled with Alamar blue and fluorescence measured at 530 nm and 590 nm. All assays were performed in triplicate.

#### Caenorhabditis elegans

Developmental and lifespan assays were conducted with N2 wild type *C. elegans* worms reared on NGM agarose plates infused with silvestrol and seeded with 30 µl OP50 E.coli/LB medium. For the developmental assays, ten worms were allowed to lay eggs for 2 h (synchronization), single eggs were isolated on separate experimental plates and incubated at 20 ℃, and the first egg laying event after 59 h was determined. Number of progeny was determined by counting the total number of progeny from synchronized, isolated animals [66]. For the lifespan assays, 40 worms were allowed to lay eggs for 2 h (synchronization), 15 eggs per plate were transferred to experimental plates and incubated at 20 ℃, and after three days, mothers were transferred to a fresh plate every other day until day eight of adulthood to avoid overgrowth by the progeny. Live worms were counted daily until all of them died. Unnatural deaths were removed from the analysis. All assays were done at least in triplicate.

#### Schistosoma mansoni

Adult worm couples were cultured in M199 medium (Sigma-Aldrich, Germany) supplemented with 10% newborn calf serum, 1% 1 M HEPES and 1% ABAM solution (10,000 units/ml penicillin, 10 mg/ml streptomycin and 25 mg/ml amphotericin B) at 37 °C in a 5% CO_2_ atmosphere. Activity of silvestrol (100 and 200 nM) against the worms was evaluated for seven days *in vitro*. Medium and silvestrol were refreshed daily and worm motility as well as the number of laid eggs assessed after three and seven days using an inverted microscope (Labovert, Germany). Worm motility was scored as recommended by WHO-TDR [67], where a score of ‘3’ indicates normal motility, ‘2’ reduced motility, ‘1’ minimal and sporadic movements, and ‘0’ represents dead worms (no movement within 30 s). Worms were obtained from infected hamsters as described elsewhere [68].

All animal experiments with Syrian hamsters (*Mesocricetus auratus*) were conducted in accordance with the European Convention for the Protection of Vertebrate Animals used for Experimental and Other Scientific Purposes (ETS No 123; revised Appendix A) and were approved by the Regional Council (Regierungspräsidium) Giessen, Germany (V54-19 c 20/15 h 02 GI 18/10 Nr. A 14/2017).

#### Leishmania amazonensis

Promastigotes of a *L. amazonensis* strain expressing *β*-lactamase [69] were cultured in immortalized J774 macrophage cells grown in RPMI supplemented with 10% of FBS and 1% PSG. Briefly, duplicate assays with silvestrol and triplicate controls were performed with plated macrophages infected with stationary phase *L. amazonensis* (25 parasites/macrophage) and incubated overnight at 32 °C and 5% CO_2_. Serial dilutions of silvestrol were added and incubated for 96 h at 32 °C and 5% CO_2_. Viability assays were conducted using CENTA™ β-Lactamase Substrate (EMD Chemicals) and Nonidet P-40 (Igepal CA 360, Fluka), and absorbance was measured at 405 nm.

#### Plasmodium falciparum

Fluorescence-based viability assays were conducted for 96 h with erythrocytic asexual cultures (5% hematocrit) of *P. falciparum* strain 3D7 (0.25% ringstage parasitemia; synchronous) in RPMI medium (RPMI 1640; 25 mM HEPES; 10 ug/ml gentamycin; 0.5 mM hypoxanthine; pH 6.75; 25 mM sodium bicarbonate; 0.5% Albumax II; 1% O_2_, 5% CO_2_: 94% N_2_) [70]. Viability was determined by quantifying fluorescence following staining of *P. falciparum* cells with SYBR Green I (Molecular Probes).

## Supplemental materials

**Table S1 – Summary of pathogens that have been previously tested for their sensitivity to a range of natural and synthetic rocaglates.**

**Table S2 – List of rocaglate-associated aa motifs extracted from a global survey of eIF4A protein sequences**

**Table S3 – List of organisms containing eIF4A isoforms with differing rocaglate-associated aa motifs**

**Table S4 – List of eIF4A mutants generated for this study Table S5 – Mutant TSA analysis**

**Table S6 – Native Aedes aegypti eIF4A temperature shift analysis**

**Tabel S7 – Arg pocket analysis**

---

**Figure S1:**
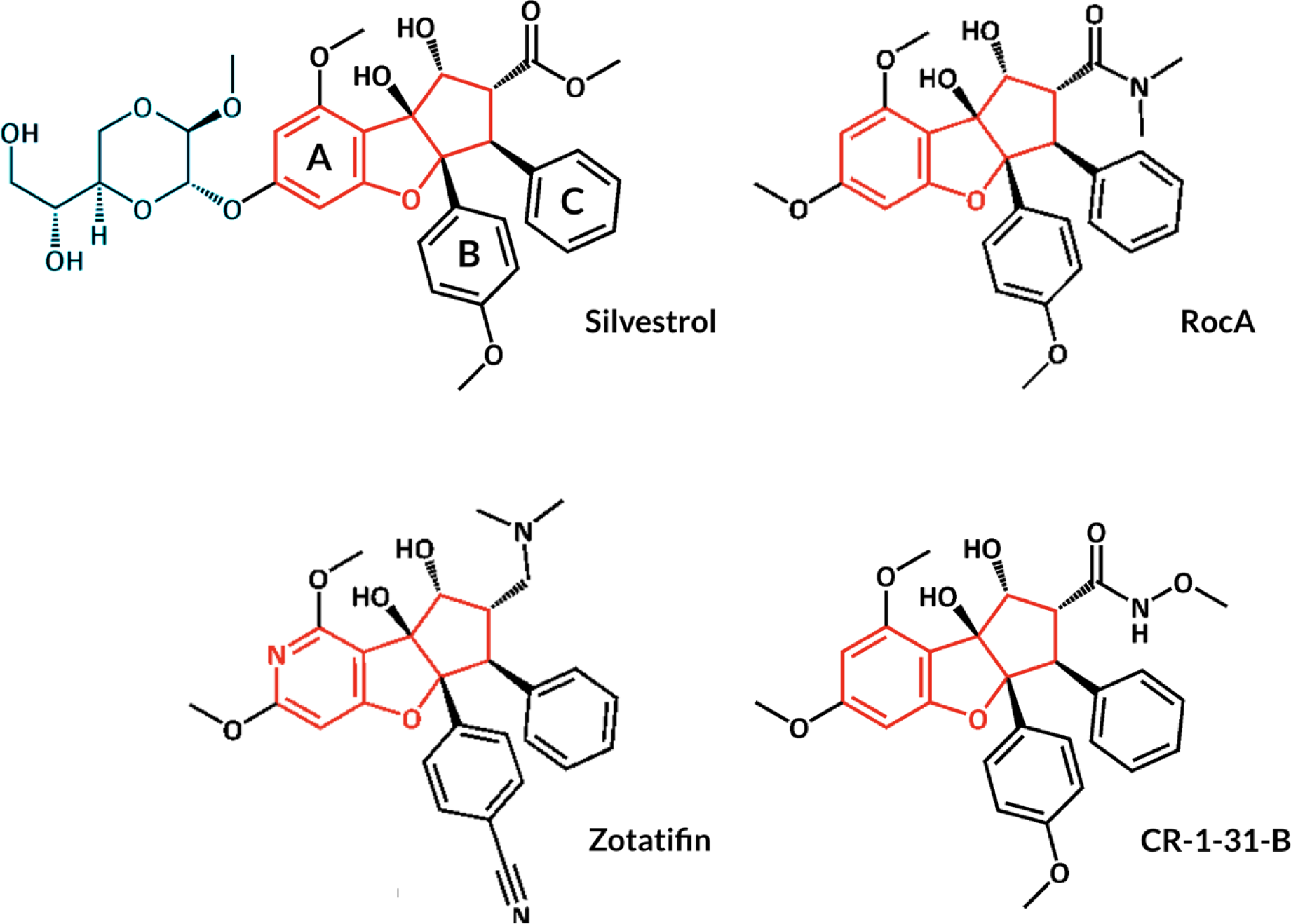
Chemical structures of the four rocaglates tested in this study. All rocaglates are characterized by a cyclopenta[*b*]benzofurane skeleton, indicated in red, and three benzyl rings, A, B, and C. Silvestrol, the archetypal natural rocaglate, contains a unique 1,4-dioxane moiety, indicated in blue, that increases the possibility of interactions with aa residues beyond those in the RNA-binding pocket. RocA, the first natural rocaglate to be purified and structurally characterized, exhibits the core structure of natural rocaglates. Synthetic rocaglates zotatifin and CR-1-31-B exhibit nitrile and imido groups, respectively, that modulate the binding characteristics of the molecules to the eIF4A:RNA complex.

**Figure S2:**
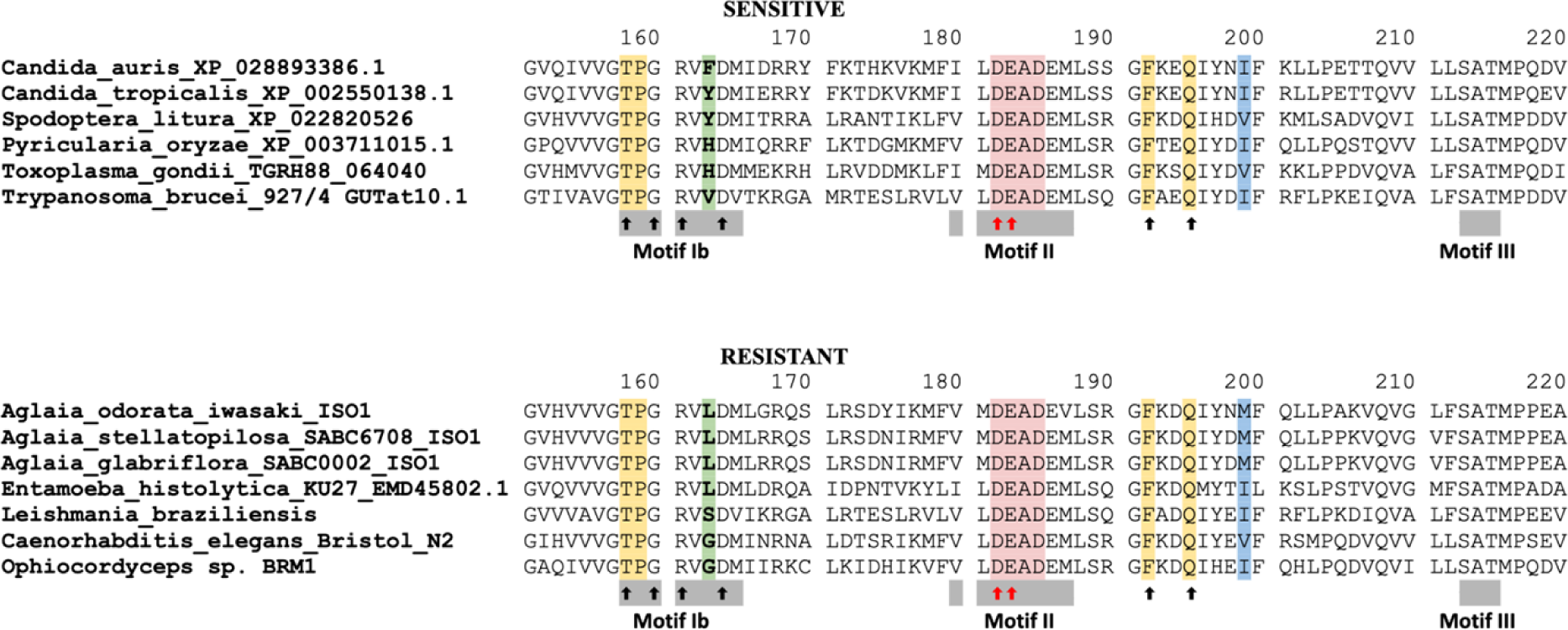
Partial aa sequence alignment of eIF4A proteins encoded by rocaglate sensitive and resistant organisms. The alignment shows the RNA-binding pocket of eIF4A, which comprises the conserved amino acid sequence Asp-Glu-Ala-Asp (D-E-A-D) characteristic of ATP dependent DEAD-box RNA helicases, motifs Ib, II, and III, which are characteristic of eIF4A proteins, and six aa residues critical to rocaglate binding. The aa residues at positions 158, 160, 161, 164, 192, and 195 (black arrows) are involved in the protein’s interaction with RNA, and residues 182 and 183 (red arrows), located within the DEAD-box (pink), are involved in the interaction with ATP. Position 163 is the primary determinant of sensitivity to rocaglates (green), followed by aa residue 199 (blue) and four conserved aa residues at positions 158, 159, 192, and 195 (yellow). The sequences shown are representative of eIF4A proteins with aa patterns that have been shown to confer rocaglate sensitivity or resistance, including two novel sequences of the rocaglate-producing plants *Aglaia stellatopilosa* and *A. glabriflora*, both reported in this study (**Accession numbers**).

**Figure S3:**
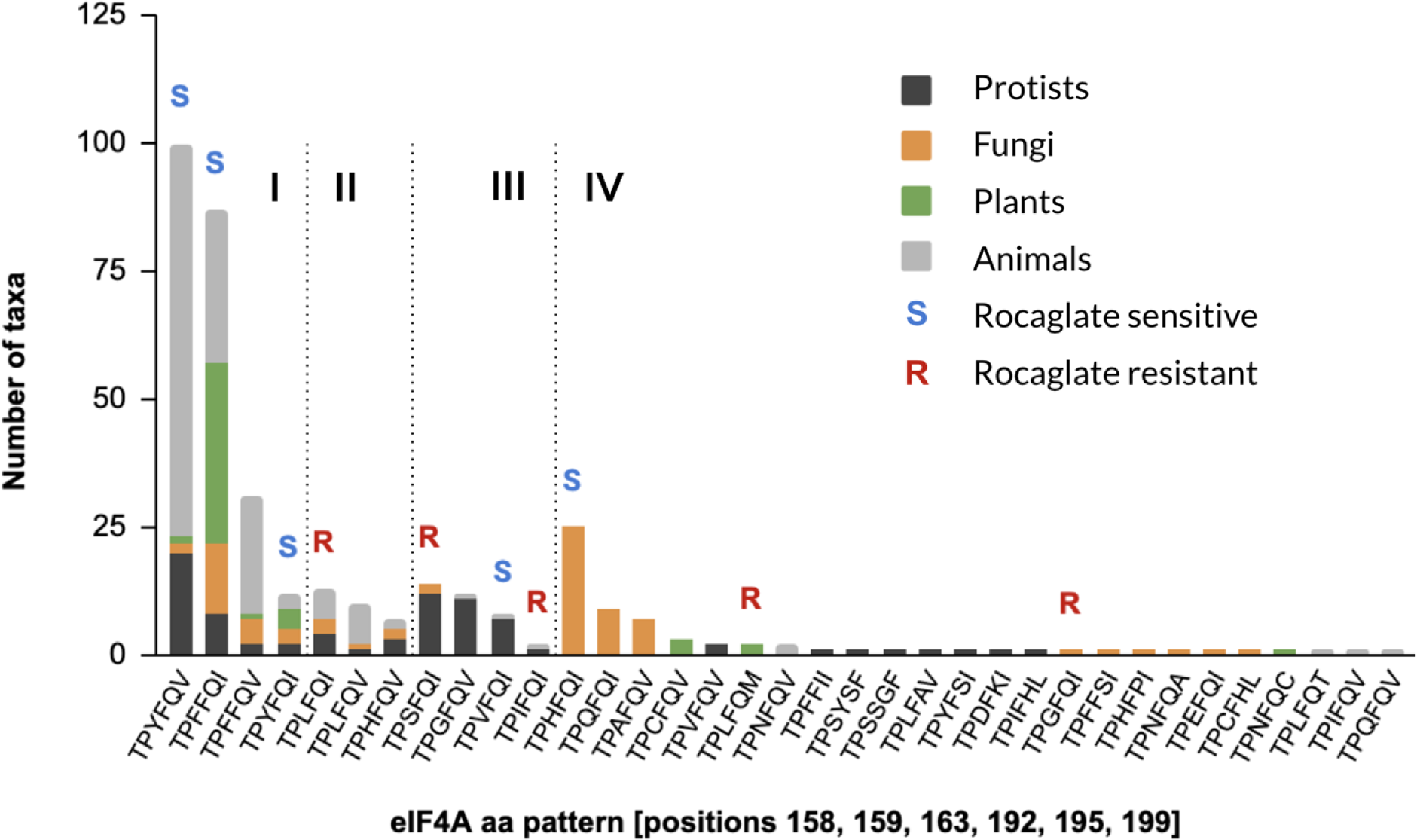
Representation of patterns of aa critical for rocaglate-binding in known eIF4A proteins across the four main groups of eukaryotes. A comprehensive analysis of known eIF4A proteins revealed 35 patterns of aa at positions 158, 159, 163, 192, 195, and 199 (human numbering). Four primary aa patterns were present in all four groups of eukaryotes, representing 63% of all eIF4As (I). Another three patterns were present in three groups (II) and four were present in two groups (III). The largest proportion of patterns, 71%, was only present in one group of eukaryotes and in most cases with only one representative species (IV). Known natural resistance is restricted to only four patterns, including two patterns—TPLFQM and TPGFQI—unique to members of the plant genus *Aglaia sp.*, so far, the only organism known to biosynthesize rocaglates, and its fungal parasite *Ophiocordyceps* sp. BRM1, respectively.

**Figure S4:**
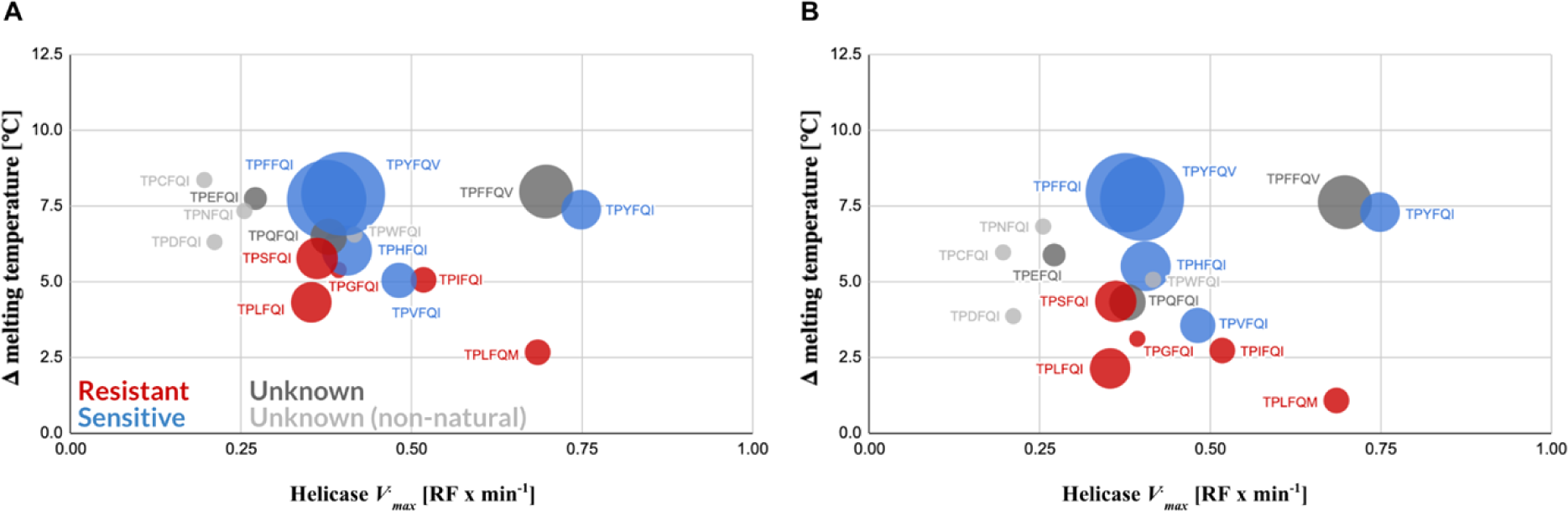
Shifts in thermal denaturation temperature of different eIF4A:RNA:zotatifin and eIF4A:RNA:CR-1-31-B complexes and their association with eIF4A sensitivity to rocaglates. The complexes established between eIF4A:RNA with two artificial rocaglates, zotatifin (**A**) and CR-1-31-B (**B**), exhibited thermal shift patterns showing a clear association between sensitivity to rocaglates and higher thermal denaturation differentials similar to those exhibited by the equivalent eIF4A:RNA:silvestrol complexes. The complexes also showed similar helicase *V_max_* ranges to those determined for the equivalent eIF4A:RNA:silvestrol complexes (**Figure 4**). Data points represent mean values of three technical replicates. Standard errors for the helicase activities are indicated in Fig. 4A and standard errors for the ***Δ*** melting temperature are listed in Table S5. The size of the circles denotes prevalence of the aa pattern among the eIF4As included in our survey.

**Figure S5:**
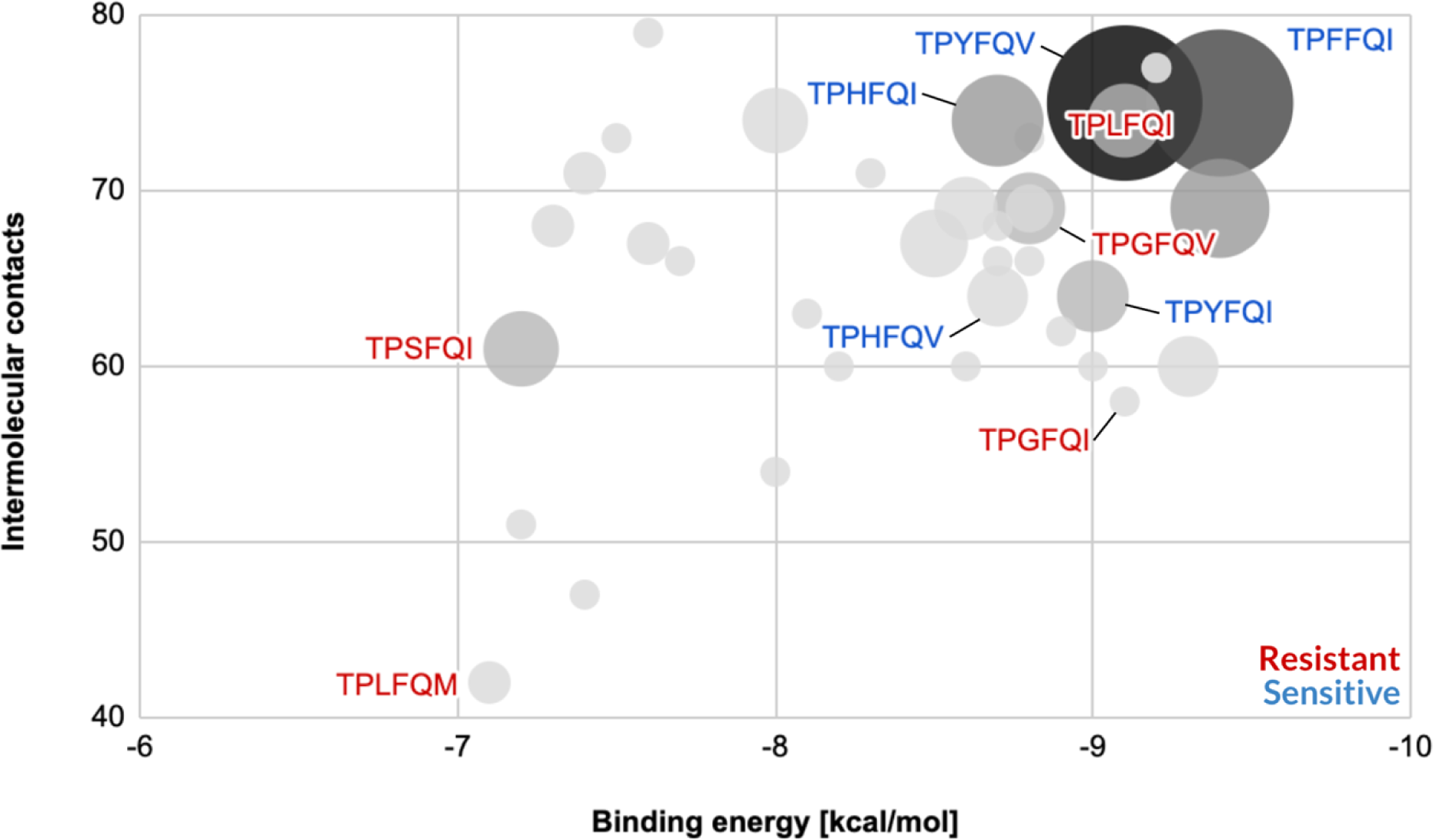
The binding energies and intermolecular contact levels of natural eIF4A:RNA:silvestrol complexes are highly correlated and exhibit an overall skew toward low binding energies and high intermolecular contacts. While rocaglate-sensitive eIF4A variants (shown in blue) exhibited the lowest binding energies and the highest intermolecular contacts, resistance to rocaglates was not preferentially associated with either of these parameters (shown in red).

**Figure S6:**
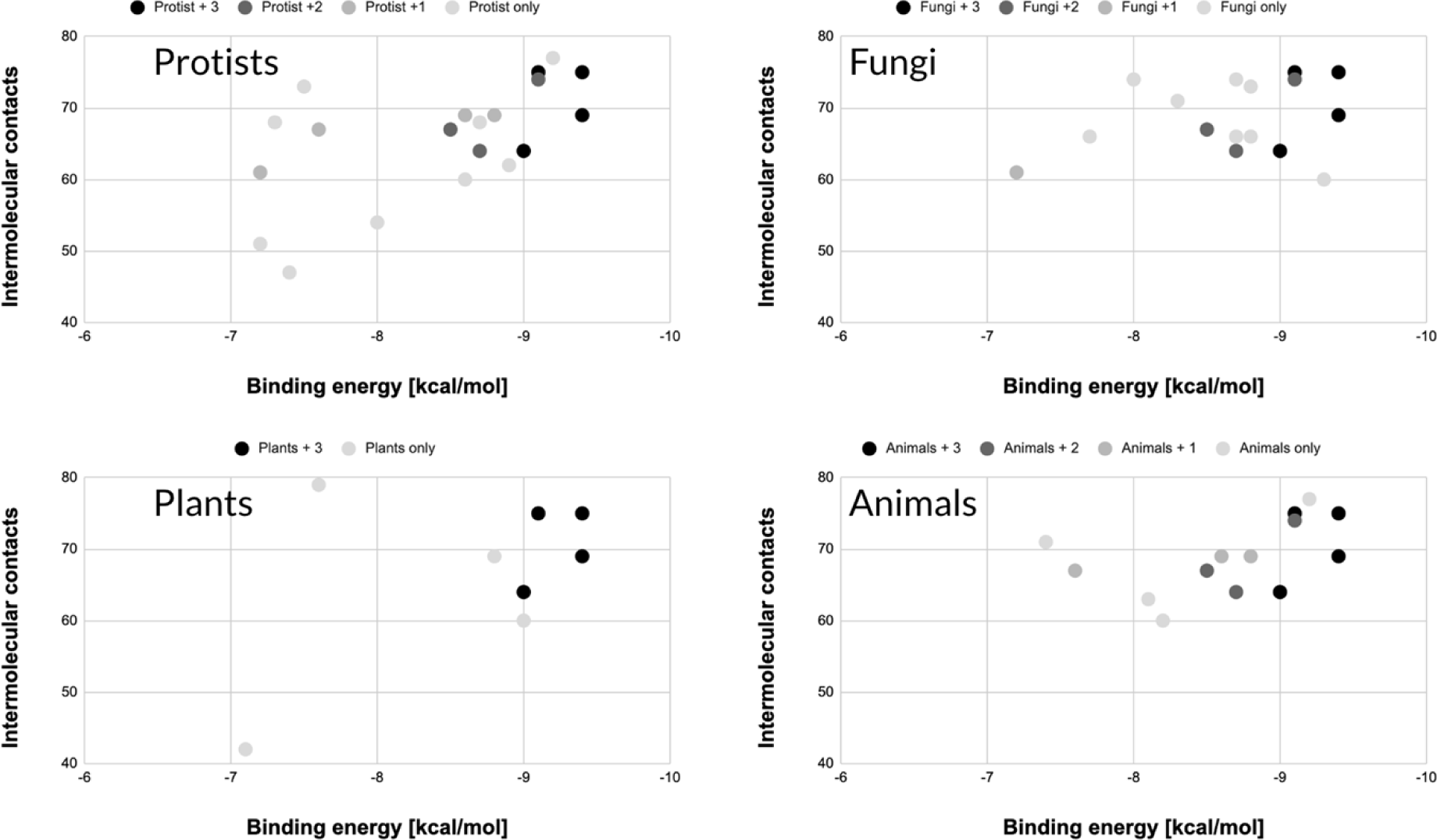
Convergence of eIF4A variants toward low binding energy/high intermolecular contact variants relative to silvestrol. In all four groups of eukaryotes analyzed—protists, fungi, plants, and animals—the inferred binding energy and intermolecular contacts converged toward variants exhibiting low binding energies and high intermolecular contact numbers. ‘+ 1’, ‘+ 2’, and ‘+ 3’ denote number of other groups of eukaryotes a particular variant is found in (e.g., “Protists + 1” denotes a variant found in protists and one of the other groups—fungi, plants, or animals).

**Figure S7:**
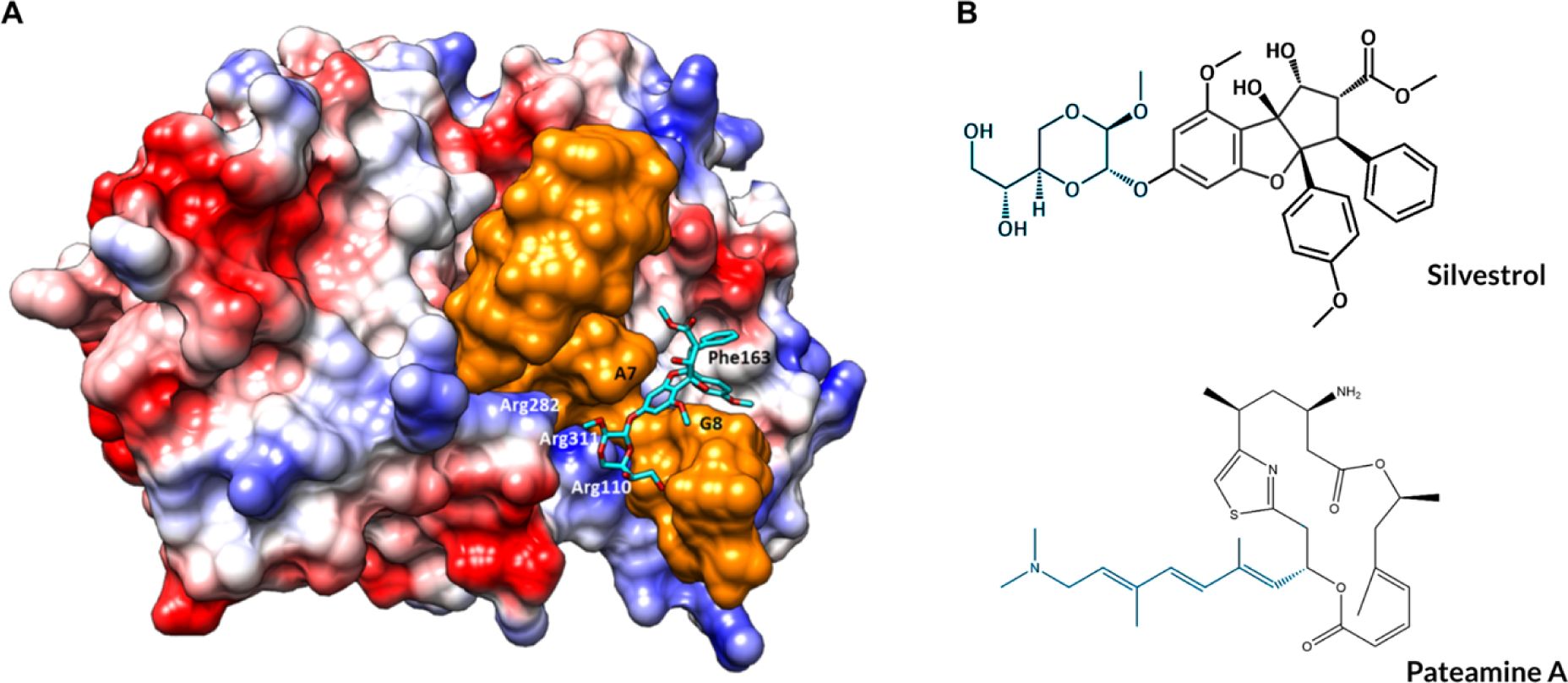
Expanded intermolecular contact interface between eIF4A and silvestrol or PatA. (**A**) Structure-based computational modeling of silvestrol onto the eIF4A-RNA complex (PDB: 5ZC9) illustrates how the 1,4-dioxane moiety of silvestrol can form additional contacts to those inside the RNA-binding pocket with three Arg residues on the surface of eIF4A— Arg110, Arg282, and Arg311—that form a highly conserved ‘Arg pocket’, resulting in a tighter clamp than the one generated by smaller rocaglates not having the 1,4-dioxane moiety. (**B**) Silvestrol and PatA exhibit analogous interactions with this Arg pocket via their 1,4-dioxane and trienyl moieties, respectively (highlighted in blue). (Electrostatic surface coloring of eIF4A generated with UCSF Chimera [blue: positively charged, red: negatively charged]).

**Figure S8:**
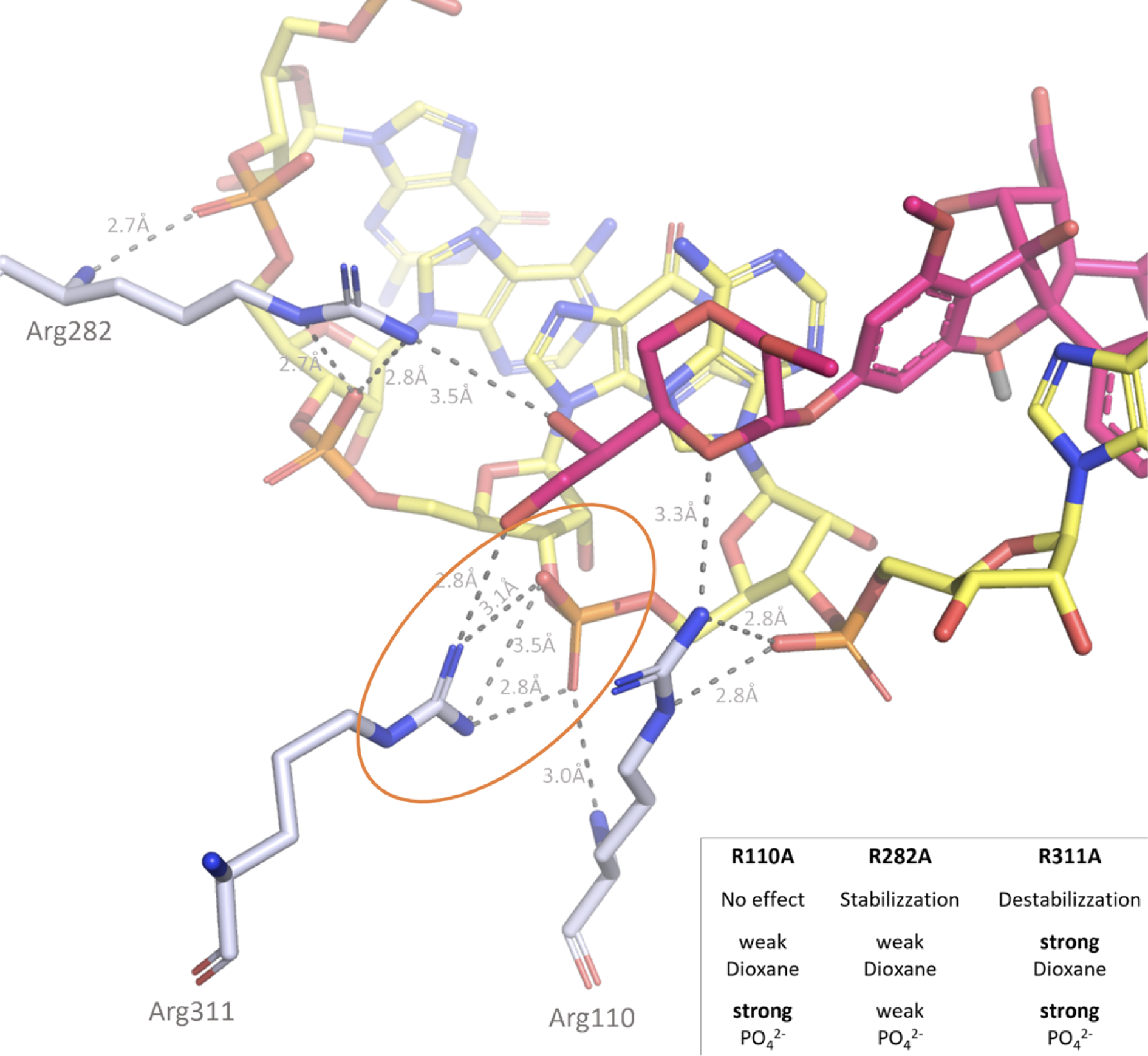
Structural interactions of the eIF4A:RNA:silvestrol complex in the Arg pocket of eIF4A. The dioxane moiety of silvestrol (magenta) interacts with the Arg rich pocket of eIF4A (grey stick) formed by Arg110, Arg282 and Arg311 (nitrogen atoms in blue and oxygen atoms in red). The Arg rich pocket is not only involved in a hydrogen bonding network (grey dashed lines) with silvestrol but it seems to play an important role also in the eIF4A-(AG)_5_ complex formation. Based on both mutation and docking studies, we suggest the order of importance of Arg residues in RNA and silvestrol binding to be Arg311 > Arg110 > Arg282. Replacing these arginine key residues by alanine, the eIF4A_(19-406)_:(AG)_5_ complex cannot be formed efficiently or at all. Normally, rocaglates increase eIF4A_(19-406)_:(AG)_5_ complex stability by raising the T*_m_* by 9.3°C (silvestrol), 8.4 °C (CR-1-31-B) and 8.0 °C (RocA). The R282A mutant shows also increased complex stability after addition of rocaglates (5,0 °C silvestrol, 3,9 °C CR-1-31-B and 3,6 °C RocA) when compared to the mutant protein alone, albeit to a lower extend (see Table S7). This may indicate that ternary complex formation is still possible but with somewhat reduced stability and is probably due to the weak interactions that Arg282 mediates with both the RNA A6 and the dioxane moiety. Therefore, this residue is important but not essential in the complex formation process. For the R110A mutant, there is virtually no difference in melting temperatures upon addition of the rocaglates (see Table S7). This suggests that rocaglates are not binding anymore to this mutant. Likely, R110 is involved in the eIF4A_(19-406)_:(AG)_5_ complex formation due to strong interaction with the phosphate group of RNA G8 which is not occurring in case of the Ala mutant. Moreover, Arg110 mediates only weak interactions with the dioxane moiety. The most important residue of the series seems to be R311. When this residue is mutated to Ala, T*_m_* is reduced upon addition of the rocaglates in a range between -3.1 and -6.3°C (see Table S7) indicating that the protein is evenly destabilized. Arg311 likely establishes a strong salt bridge (orange circle) with the phosphate groups of RNA A7 (yellow sticks with phosphorus atoms colored orange) and in a strong hydrogen bond with the dioxane moiety of silvestrol. An R311A mutant precludes the formation of a stable eIF4A-RNA complex in the presence of rocaglates (see **inset**). Most likely, the equilibrium between complex formation and dissociation shifts toward dissociation because of the loss of the strong salt bridge with the RNA. Under these conditions, rocaglates cannot bind as well because the complex is probably not in the most favorable conformation to allow binding to occur. This is potentially the limiting step of complex formation: only when the RNA binds to eIF4A in the optimal conformation, the inhibitors can clamp and bind to the eIF4A-(AG)_5_ complex, otherwise they probably bind more loosely, and they even destabilized the whole system. The behavior of the triple mutant is comparable with the single mutant R311A. Although no synergistic effect of the three mutated Arg residues is noted, a destabilizing effect with T*_m_* reduction ranging from -3.2 °C to -3.6 °C is observed (see Table S7).

**Figure S9:**
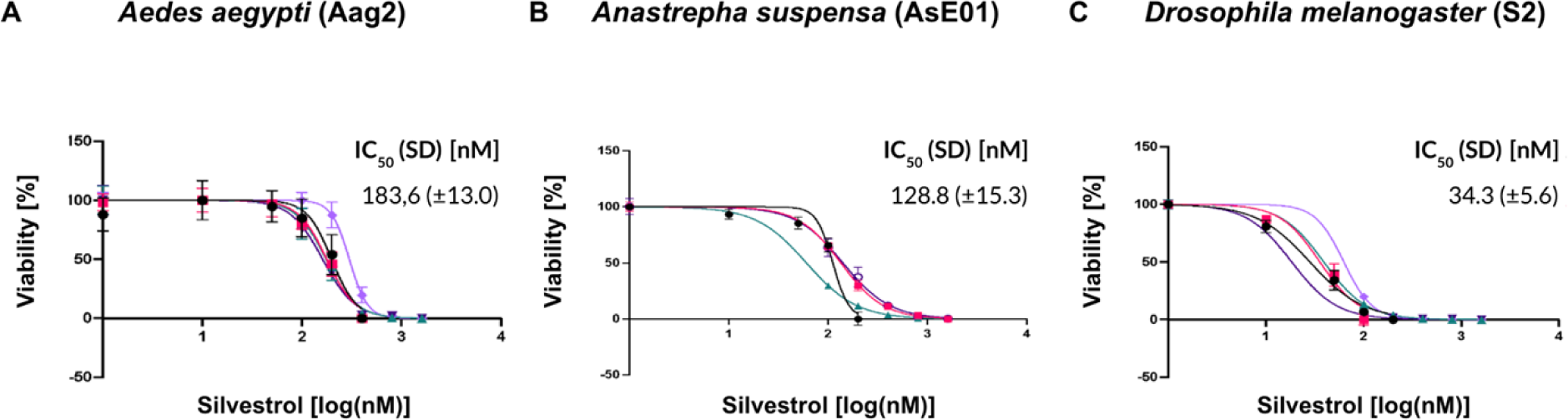
*In vitro* assays of sensitivity to silvestrol in mosquito and fruit fly cell lines expressing eIF4A proteins with previously untested and tested rocaglate-associated aa patterns. In all instances, viability was measured using a cell proliferation assay. (**A**) *Aedes aegypti* (Aag2) [TPHFQV; sensitive], (**B**) *Anastrepha suspensa* (AsE01) [presumably TPYFQV; sensitive], (**C**) *Drosophila melanogaster* (S2) [TPYFQV; sensitive].

**Figure S10:**
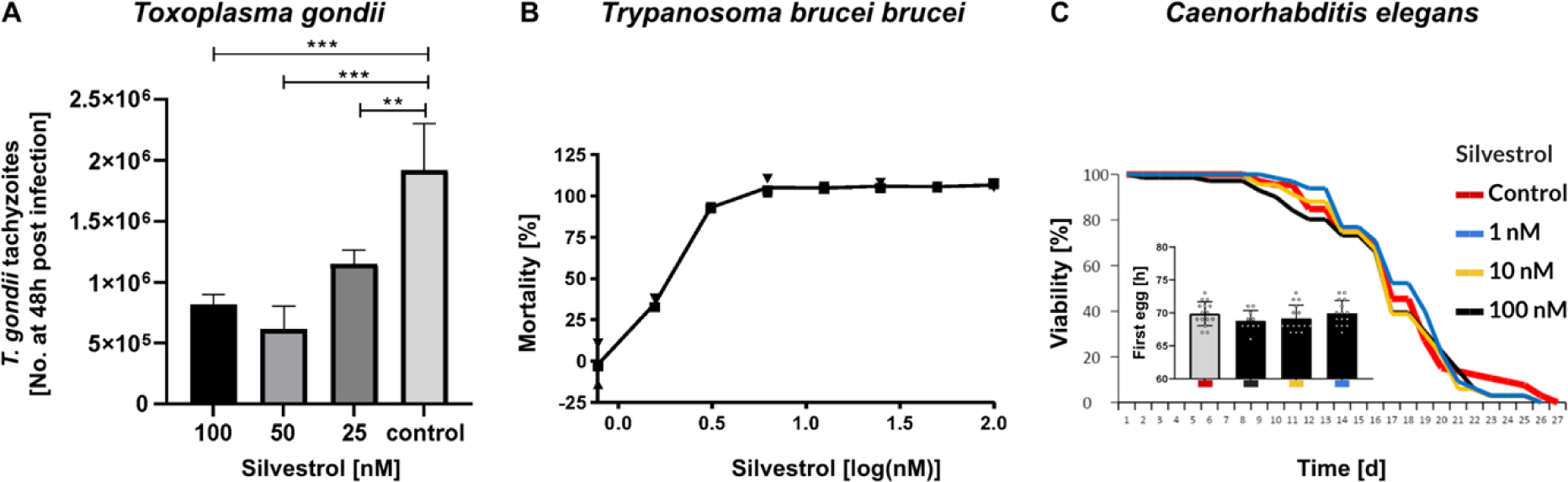
*In vitro* assays of sensitivity to silvestrol in organisms expressing eIF4A proteins with previously untested rocaglate-associated aa patterns. (**A**) *Toxoplasma gondii* [TPHFQV; sensitive] replication was measured using MARC145 monkey kidney cells as the host. (**B**) *Trypanosoma brucei brucei* [TPVFQI; sensitive] mortality was determined using a free parasite viability assay. (**C**) *Caenorhabditis elegans* [TPGFQV; resistant] viability and developmental pace (inset) were measured on plates.

**Figure S11:**
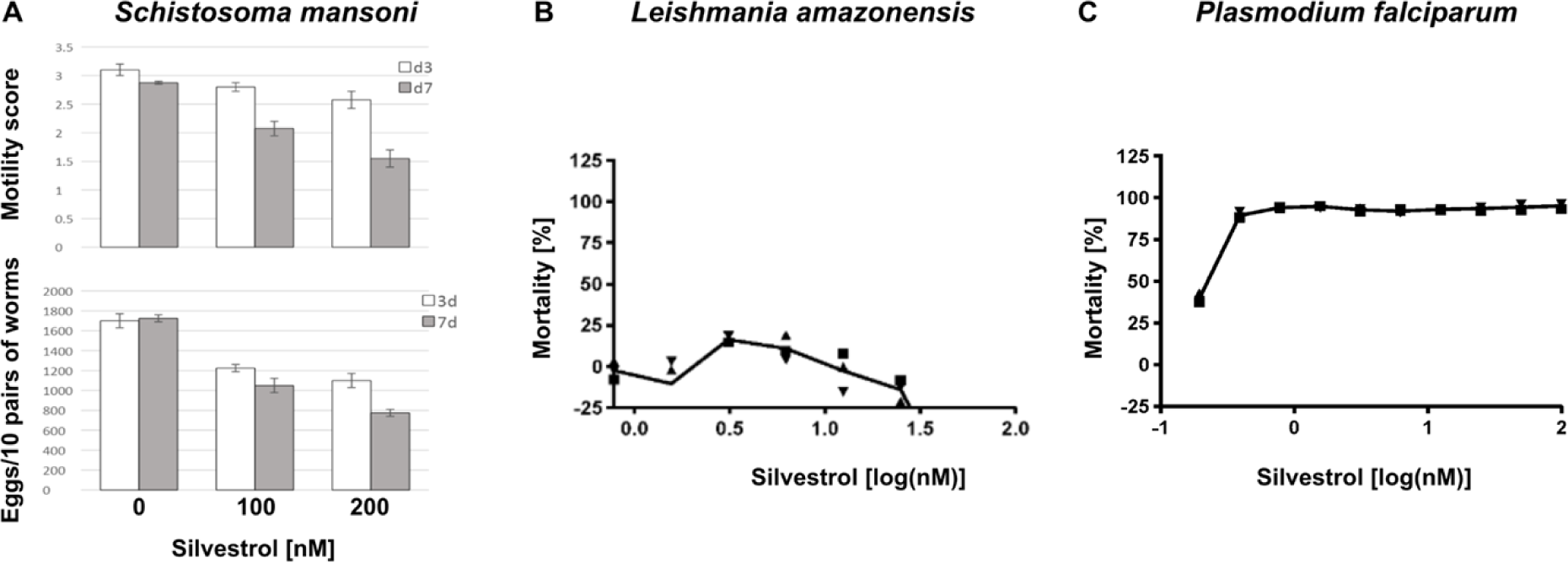
*In vitro* assays of sensitivity to silvestrol in organisms expressing eIF4A proteins with previously tested rocaglate-associated aa patterns. (**A**) *Toxoplasma gondii* [TPHFQV; sensitive] replication was measured using MARC145 monkey kidney cells as the host. (**B**) *Trypanosoma brucei brucei* [TPVFQI; sensitive] mortality was determined using a free parasite viability assay. (**C**) *Caenorhabditis elegans* [TPGFQV; resistant] viability and developmental pace (inset) were measured on plates.

## Acknowledgments

The authors would like to thank Christina Scheld, Georgette Stovall, and Christin Ritter for excellent technical assistance.

## Author Contributions

Conceptualization: GTO, AG & TCY

Data curation: WO, MFDBA, LK, NS, FM, JS, LMRS, CH, AHL, HH, SH, IH, GL & GTO

Formal analysis: WO, MFDBA, AG & GTO

Funding acquisition: MFS, CGG, FCS, AT, AR, AH, TCY & AG

Investigation: WO, MFDBA, LK, NS, FM, JS, LMRS, AHL, HH, SH, MYL & IH

Methodology: WO, MFDBA, JS, FM, AHL & SH

Project administration: AG, TCY & GTO

Supervision: WO, GL, TCY, AG & GTO

Visualization: WO, JS, AHL, SH & GTO

Writing – original draft: GTO

Writing – review & editing: WO, MFDBA, LK, NS, FM, JS, LMRS, CH, AHL, HH, SH, IH, GL, MFS, CGG, FCS, AT, AR, AH, TCY & AG

### Funding

This work was funded by the LOEWE Center DRUID (project A2 to AG, A4 to AH, B4 to CGG, D4 to CH and AT, D5 to MFS and IH, and E1 to SH and CGG) and the German Ministry of Science and Education (BMBF) via the HELIATAR project (project 16GW0259 to AH and project 16GW0258K to AG) and the DLR project (project 01DG20023 to CH)). FCS and AHL’s research was supported in part by the National Institutes of Health (R35GM131877 and U01GM110714 to FCS) and the Howard Hughes Medical Institute (Faculty Scholar Grant to FCS).

